# Oscillatory Co-expression of HES1 and HES5 Enables a Hybrid State in a Bistable Transcription Factor Regulatory Motif

**DOI:** 10.1101/2025.05.22.655498

**Authors:** Veronica Biga, Anzy Miller, Anoushka Kamath, Ying Q P Mak, Antony D Adamson, Elli Marinopoulou, Paul François, Nancy Papalopulu, Cerys S Manning

## Abstract

Many cell fate decisions in the developing neural tube are directed by cross-repressive transcription factor (TF) motifs that generate bistability, enforcing expression of one dominant TF. However, evidence of hybrid states, where cells co-express opposing fate determinants, challenges this model. We hypothesised that oscillatory expression enables co-existence of cross-repressive TFs within single cells, allowing hybrid states in bistable motifs. To test this, we focused on HES1 and HES5, oscillatory, cross-repressive TFs that regulate neural progenitor maintenance and are expressed in adjacent dorsoventral domains in the developing spinal cord. Using live-cell imaging of fluorescent reporters and computational modelling, we show that HES1 and HES5 co-express and oscillate in-phase within single cells. Differences in protein stability result in distinct free-running periodicity, but co-expression results in entrainment and phase-locking. Modulating cross-repression strength and/or abundance shifts the system towards bistability and dominance of a single TF oscillator. Consistent with this, we observe progressive separation of the HES expression domains in vivo, through a decrease in oscillatory co-expression. Our findings provide a mechanism for hybrid states to emerge in a developmental bistable motif.

## Introduction

In the developing neural tube, progenitor cells make decisions to differentiate into specialised cell types, governed by gene expression programmes that define specific cell fates. Along the dorsoventral axis of the mammalian spinal cord, the complexity of cell types arises from progenitor domains which are reproducibly organised in distinct domains of gene expression. These arise when homeodomain transcription factors (TF), such as Pax6, Olig2 and Nkx2.2, are differentially induced by morphogenetic factors and then, cross-repress each other in adjacent progenitor domains, reinforcing distinct gene expression territories (Briscoe et al., 2000, Balaskas et al., 2012, Sagner and Briscoe, 2019). Domain boundaries, which initially contain cells of different fates (expressing different genes) are progressively sharpened, ensuring stable, region-specific cell fates along the dorsoventral axis. These domain-specific expressions of TFs, reflecting mutually exclusive cell fates, reflect several bistable transcriptional switches between TFs in adjacent domains, which together comprise an extensive gene regulatory network. In this case, bistability, a system in which cells can stably adopt one of two gene expression states, is mediated by cross-repression between fate-determining TFs, where cells can express gene A or B but not both (Ferrell, 2002). Bistability enables cells to switch between discrete states and is a common mechanism for enforcing binary fate decisions in development (Moris et al., 2016, Kutejova et al., 2016).

In traditional or idealised models of bistability, intermediate states in which cells express both gene A and B are unstable with abrupt transitions between A+/B- and A-/B+ states (Wang et al., 2009, Verdugo et al., 2013, Zorzan, 2021, Rombouts and Gelens, 2021). However, during development there is growing evidence to suggest that before fates are resolved, progenitor cells can occupy a transitory or “hybrid” state, in which genes associated with opposing fates are co-expressed in the same cells, as discussed in (Moris et al., 2016). For example, in early embryonic development co-expression of pluripotency markers (e.g OCT4) and TFs associated with lineage commitment such as GATA6, MIXL1, Hex as well as PDGFRa have been reported (Plusa et al., 2008, Morgani et al., 2013, Allison et al., 2018, Stavish et al., 2020, Redo-Riveiro et al., 2024). In the developing nervous system, we have detected co-expression of progenitor and differentiation genes by smFISH the zebrafish hindbrain (Soto et al., 2020). scRNAseq is also increasingly identifying hybrid states in many different contexts where cell fate decisions are made, including hematopoiesis (Hu et al., 1997, Laslo et al., 2006, Olsson et al., 2016, Bergiers et al., 2018), osteogenesis (Wang et al., 2009), T cell differentiation (Zhou et al., 2008). The abundant detection of intermediate or hybrid states in scRNAseq across different biological contexts, reviewed in (MacLean et al., 2018) has also spurred new computational approaches to determine the underlying network structure (Dey, 2025) as well as identify such states from data, for example (Kong et al., 2022).

This presents a conceptual challenge: how can cross-repressing TFs coexist within the same cell, and how is this hybrid state eventually resolved into a stable mutually exclusive fate choice? A few theoretical explanations have been proposed, including narrow parameter regimes which permit co-expression (Andrecut et al., 2011), weak (Laslo et al., 2006, Rohm, 2018) or delayed cross-repression (Zhu et al., 2007, Bokes et al., 2009) and noise from transcriptional bursting (Bokes et al., 2013). In theoretical models of spinal cord patterning, the network structure and dynamics of cross-repressive switches allow for hybrid states to be observed transiently (Perez-Carrasco et al., 2016, Exelby et al., 2021). Further, modelling has suggested that oscillations can lead to co-expression in a bistable switch comprised of competitive TFs (Bokes and King, 2019). However, experimental evidence is lacking. We hypothesised that oscillatory expression of TFs within the same cell may allow progenitor cells to co-express cross-repressive fate determinants, offering a mechanistic basis for the observed hybrid state, while at the same time allowing the fates to be resolved over time. To test this hypothesis in the context of spinal cord development, and to gain some mechanistic insight, we focused on HES1 and HES5, members of the basic helix-loop-helix (bHLH) family of transcriptional repressors, because they are known to have both cross-repressive interactions, and oscillatory expression during neural development. Evidence for cross-repression comes from single knock-out mouse models, where loss of *Hes1* leads to *Hes5* becoming expressed throughout spinal cord; conversely loss of *Hes5* causes *Hes1* to be upregulated in areas where *Hes5* would normally be highly expressed (Hatakeyama et al., 2004). Furthermore, over-expression of Hes5 in mouse embryos leads to a decrease in Hes1 expression and conversely, Hes5 is upregulated in Hes1 germline mutants (Bansod et al., 2017, Riesenberg et al., 2018).

The Notch target genes HES1 (including zebrafish ortholog Her6) and HES5 have been shown to have temporal oscillatory expression across different species and tissue types including the mouse spinal cord (Manning et al., 2019, Biga et al., 2021, Hawley et al., 2022), forebrain (Imayoshi et al., 2013, Ochi et al., 2020), pre-somitic mesoderm (Niwa et al., 2007, Shimojo et al., 2016, El Azhar et al., 2024) and the zebrafish brain (Soto et al., 2020, Doostdar et al., 2024). HES oscillations are typically in the order of a few hours (Kageyama et al., 2007, Manning et al., 2019, Soto et al., 2020, Doostdar et al., 2024) and are generated via auto-negative feedback (Takebayashi et al., 1994, Kageyama et al., 2007) combined with short mRNA and protein half-lives (Hirata et al., 2002, Imayoshi et al., 2013). Double HES knock-outs show that their expression is required for central nervous system develop and important for neural progenitor maintenance (Hatakeyama et al., 2004, Bansod et al., 2017) while experiments involving characterisation and manipulation of dynamics shows that their oscillatory expression is important for cell state transitions (Manning et al., 2019, Soto et al., 2020, Marinopoulou et al., 2021, Maeda et al., 2023, Shimojo et al., 2024) .This indicates that oscillatory HES represent a developmental program responsible for enabling state transitions in the developing CNS including spinal cord. Much like the homeodomain TFs mentioned above, HES1 and HES5 are thought to occupy mutually exclusive adjacent territories along the dorsoventral axis of the developing spinal cord (Sagner et al., 2018), which combined with single HES mouse knock-out data, suggest that this is due to cross-repression (Hatakeyama et al., 2004). However, it was not known whether these 2 genes show simultaneous oscillatory expression in the same cells and whether the oscillators interact with each other in a switch-like way that could explain their resolution to distinct domains of expression.

To answer these questions, in this study, we performed simultaneous live imaging of HES1 and HES5 in single cells, using genetically engineered reporters for endogenous expression dynamics. We observed co-expression of HES1 and HES5 in neural stem cells in vitro, which we verified in vivo, particularly during early developmental stages. Computational modelling based on auto-and cross-repression predicted that HES1 and HES5 have different free running periods, explained in silico by differential protein stability and validated in singly HES1 or HES5 expressing cultured neural stem cells. However, when HES1 and HES5 were co-expressed in the same cells, the two oscillators became in-phase, suggesting an entrainment event whereby the slower periodicity TF (HES5) entrains the faster one (HES1). Changing the cross-repression strength or the level of either protein in silico, allowed one oscillator to prevail and switch off the other, which is likely to underlie the resolution of expression into mutually exclusive domains during development of the spinal cord. Our findings show that the simultaneous, in-phase oscillatory expression of HES1 and HES5 allows progenitor cells to co-express cross-repressive fate determinants, before resolving to mutually exclusive expression, offering a mechanistic basis for a hybrid state. Furthermore, our data provide a mechanistic link between oscillatory expression and fate resolution, and reveal additional features of HES oscillations, including entrainment and the extension of the decision-making window, potentially enhancing both robustness and flexibility in neural development.

## Results

### In vitro neural progenitor cultures indicate HES1 is co-expressed with HES5

The HES1 and HES5 TFs are critical for spinal cord development and together with their upstream Notch signalling input represent a toggle-switch, a motif known to exhibit bistability (Ferrell, 2002) **(Figure 1A)**. We defined the cell states associated with their endogenous expression using previously described knock-in fluorescent reporters (Imayoshi et al., 2013, Marinopoulou et al., 2021) in both primary neural progenitor cells (priNPCs) derived from E10.5 spinal cord (Venus::HES5^+/-^, HES1::mScarlet-I^+/-^) and mouse embryonic stem cell (mES) -derived dorsal neural progenitor cells (NPCs) (Venus::HES5^+/+^, HES1::mScarlet-I^+/+^) (**Figure 1B-G**). In both primary and mES-derived NPCs, we observed the presence of single expressing HES1-/HES5+ or HES1+/HES5-as well as a large proportion of HES1+/HES5+ co-expressing cells (**Figure B,C**). As expected, priNPC cultures express the progenitor marker SOX2 (Graham et al., 2003) (**Supplementary Figure 1A**). We quantified the proportions of DAPI-stained nuclei that co-express HES1+/HES5+ to be approx. 65% of priNPCs (**Figure 1D**). The percentage of double positives was higher (approx. 85%) in mES-derived NPCs where nuclei were detected with a PAX3/7 (Mansouri and Gruss, 1998) antibody to mark neural progenitors and exclude other cell types **(Figure 1C,E**). Overall, double positives represented the highest fraction in both culture systems (**Figure 1D,E)**. We also observed a differential bias for single expressing cells in the different cultures. Specifically, priNPC cultures contained more HES1-/HES5+ cells whereas mES-derived NPCs contained more HES1+/HES5-cells (**Figure 1D vs E**). These observations are consistent with the fact that the mES differentiation protocol enriches for dorsal types (Gupta et al., 2022) (identified by expression of PAX3/7) where HES1 is more widely expressed compared to ventral (Sagner et al., 2018); in contrast, the priNPCs could be ventral or dorsal, with HES5 being expressed in both areas (Manning et al., 2019).

**Figure 1.**
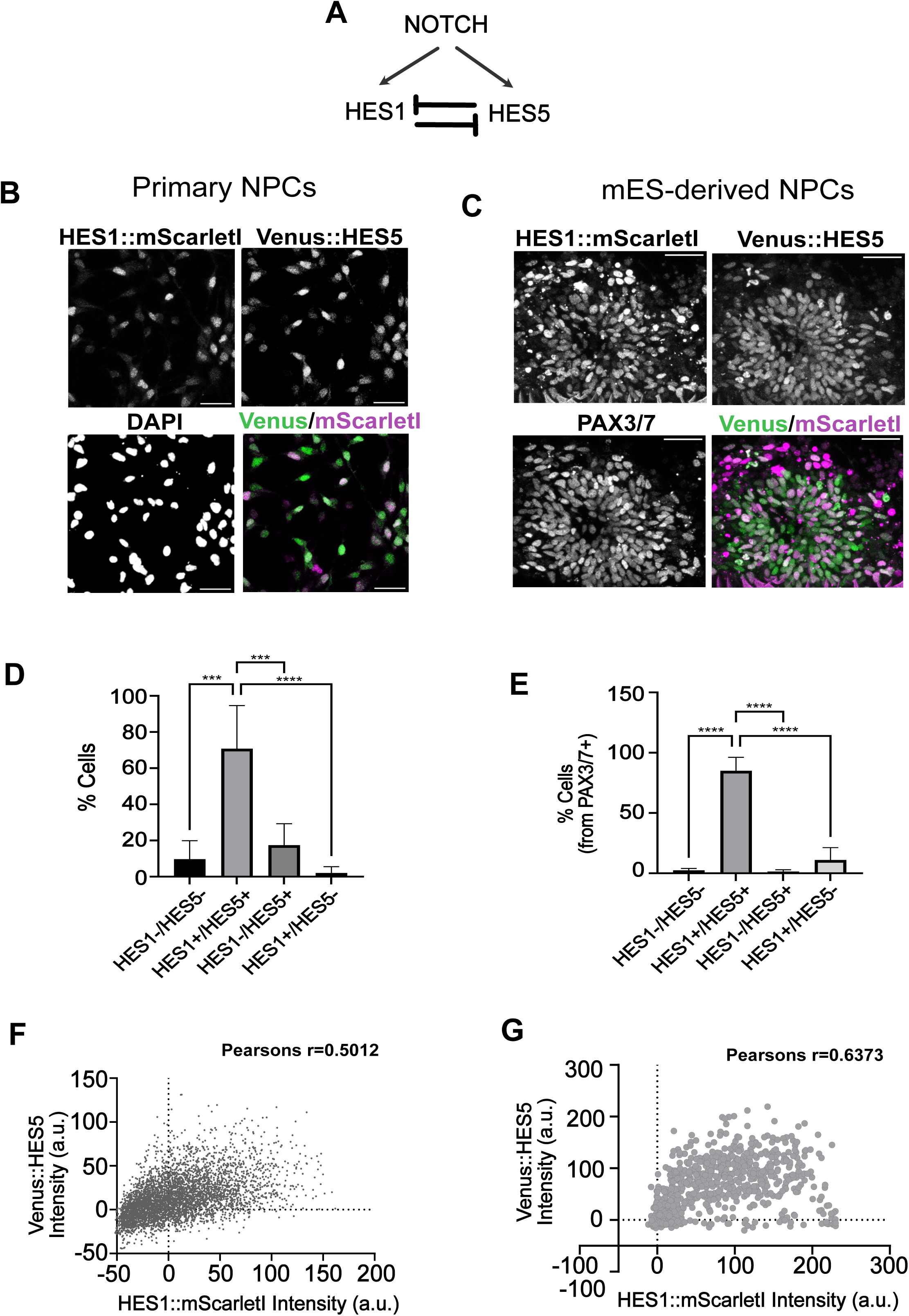
Co-expression of HES1 and HES5 in primary spinal cord and mouse embryonic stem cell (mES)-derived neural progenitor cells (NPCs). **(A)** Diagram of Notch activating the HES1 and HES5 targets which cross-repress. **(B)** Fixed primary neural progenitor cell cultures expressing endogenous HES1::mScarlet-I, endogenous Venus::HES5 and a nuclear DAPI stain at 2 days in culture; 40x objective, scale bar 40um. **(C)** Fixed mES-derived dorsal progenitor cell cultures at 2 days post-retinoic acid removal (Materials and methods) expressing endogenous HES1::mScarlet-I, endogenous mVenus::HES5 and immunofluorescence for PAX3/7 antibody; 40x objective, scale bar 40um. **(D-E)** Quantification of single and double positive HES1::mScarlet-I and Venus::HES5 fractions observed in priNPC **(D)** and mES-derived NPC **(E)**; bars indicate mean and SD; D: 4 independent experiments, >2,000 cells analysed; E: 4 neural rosettes with a total of 698 cells analysed; 1-way ANOVA, Tuckey’s multiple comparison correction with p<0.0001. **(F-G)** Scatter plot of HES1::mScarlet-I versus Venus::HES5 in the same cell obtained from priNPC **(F)** mES-derived NPC cultures **(G)**; markers indicate background-subtracted mean fluorescent intensity per nucleus from 1 independent experiment from each culture system.

We quantified the level of expression of HES1 and HES5 in the same nuclei across the two culture systems and observed positive correlation coefficient values indicative of co-expression at a range of levels (**Figure 1F,G**). These results indicated that a ‘hybrid state’ exhibiting co-expression of HES1 and HES5 is observed in most spinal cord neural progenitor cells in-vitro.

### HES1 and HES5 oscillate at a similar period although HES1 is more oscillatory and has a higher amplitude compared to HES5

Separate oscillations of HES1 and HES5 have been previously reported in primary neural progenitor cultures (Imayoshi et al., 2013, Manning et al., 2019, Marinopoulou et al., 2021), however they have not been simultaneously monitored in the same system using dual reporters. Here, we directly compare the oscillatory parameters of HES1 and HES5 in NPCs using timelapse imaging and tracking of individual nuclei (**Figure 2A-I**). We observed that both proteins have peaks of expression over time (**Figure 2A, B**- examples from 2 progenitors) with HES1 expression dynamics being more regular and periodic compared to HES5 (**Figure 2C**-log likelihood ratio, LLR indicative of oscillation quality, see (Phillips et al., 2017)). A higher proportion of HES1 timeseries passed a statistical test for oscillatory expression compared to HES5 timeseries (**Figure 2D**). In addition, the dynamics of HES1 expression were more pronounced compared to HES5, quantified as the ratio of intensity at the peak versus intensity at the trough (**Figure 2E**, peak-to-trough fold change). As the two proteins were monitored with different fluorophore fusions (HES1 fused to mScarlet-I and HES5 fused to Venus), we separately validated that this effect is not due to the different fluorophores by comparing HES1 oscillations in priNPCs containing HES1 fused on one allele with mVenus and the other allele with mScarlet-I (**Supplementary Figure 1B**, Materials and Methods). As expected, predominant periodic activity was detected in cells containing dual HES1 fluorophore fusions (**Supplementary Figure 1C**,D) and we observed no differences in the maximum or mean fold-change in these cells (**Supplementary Figure 1E**). This argued against artefactual differences due to the choice of fluorophores and confirmed that HES1 is more oscillatory and has a higher fold-change compared to HES5 in priNPCs (**Figure 2D,E**). The period of oscillations was variable at single cell level, averaging across repeats at 3.3h for HES1 and 3h for HES5 in priNPCs with no significant differences at the population level (**Figure 2F**).

**Figure 2.**
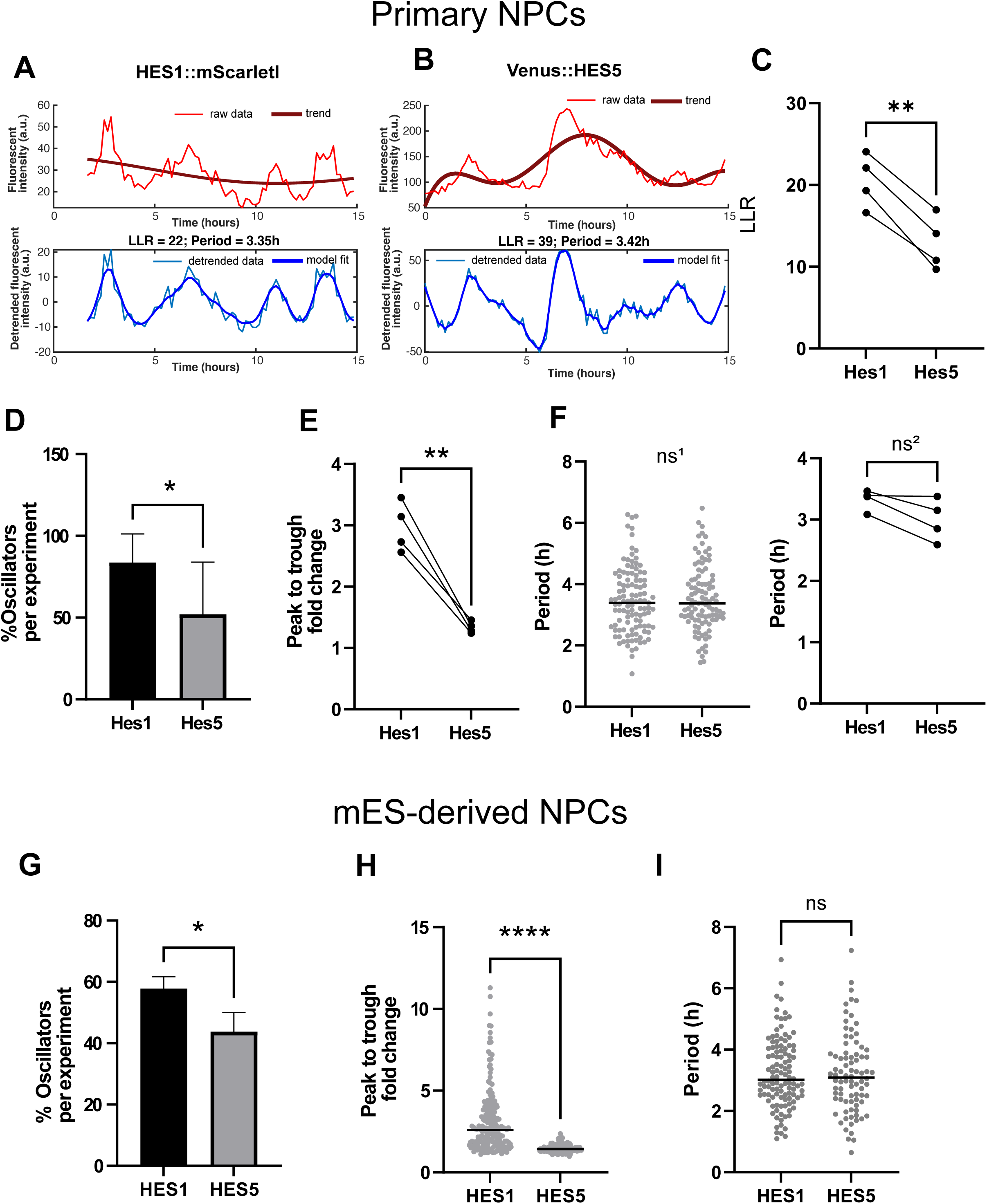
Oscillatory expression of HES1 and HES5 in primary spinal cord and mouse ES-derived neural progenitor cells. **(A-B)** Representative examples of periodic fluctuations observed in mean nuclear intensity of HES1::mScarlet-I **(A)** and Venus::HES5 **(B)** in two different priNPCs over time; upper panels indicate raw intensity and slow-varying trend (solid red line); lower panels indicate detrended intensity and periodic model fit (solid blue line); LLR denotes log-likelihood ratio. **(C)** Comparison of LLR values (indicative of oscillation quality) observed in HES1::mScarlet-I versus Venus::HES5 timeseries; markers indicate median per experiment. **(D)** Percentage of oscillatory timeseries observed in HES1::mScarlet-I and Venus::HES5; bars indicate mean and SD.**(E)** Comparison of maximum peak-to-trough fold change in fluorescent intensity per cell (Materials and methods) in HES1::mScarlet-I versus Venus::HES5 timeseries; markers indicate median per experiment. **(C-E)** paired t-test, 2 -tailed with p<0.01; sample size: 4 independent replicates with 446 tracks in total. **(F)** Comparison of period duration in oscillatory HES1::mScarlet-I versus Venus::HES5 timeseries observed in a single experiment as well as across multiple repeats; Left: markers indicate cells and bars indicate median; Mann-Whitney test, 2-tailed, non-significant ns^1^=0.7065; Right: markers indicate median per experiment; paired t-test, 2-tailed, non-significant p=0.0639. **(G)** Percentage of oscillatory timeseries observed in HES1::mScarlet-I and mVenus::HES5 observed in mES-derived NPCs; bars indicate mean and SD; paired t-test, 2-tailed with p<0.05; sample size of **(G-I)** is 3 independent experiments with 193 tracks in total. **(H)** Comparison of maximum peak-to-trough fold change in mES-derived progenitors; markers indicate cells and bars indicate median; Mann-Whitney test, 2-tailed with p<0.0001. **(I)** Comparison of period duration in oscillatory HES1-mScarlet-I versus mVenus::HES5 timeseries in mES-derived progenitor cultures; markers indicate cells, bars indicate median; Mann-Whitney test, 2-tailed, non-significant ns=0.7443.

In mES-derived NPCs we observed features of the dynamics of HES1 and HES5 expression consistent with the priNPCs (**Figure 2G-I**). The percent of time-series that pass as oscillators of either HES appeared somewhat reduced in mES-derived cultures compared to priNPCs which suggests closer similarity to tissue, such as reported for HES5 in ex-vivo mouse spinal cord (Manning et al., 2019). Like priNPCs, in mES-derived NPCs HES1 oscillations were more frequent, and had higher fold-change intensity ratios between peaks and troughs compared to HES5 oscillations. For both HES1 and HES5 expression, there is variability in the period but with no significant differences between the two (**Figure 2I**). Overall, we concluded that HES1 is more oscillatory, has a higher peak-to-trough fold change, but, on average has the same period as HES5 at single cell level.

### HES1 and HES5 oscillate in-phase in the co-expressing fraction

Having determined that HES1and HES5 can oscillate in the same culture conditions, we next investigated if their dynamics are in-phase or out-of-phase in the same cells. To do this, we monitored the fluctuations in level of both Venus::HES5 and HES1::mScarlet-I in the same cells over time (**Figure 3A-F**). The majority of HES1 and HES5 co-expressed and showed positively correlated dynamics at the single cell level in priNPCs (**Figure 3A, Supplementary Figure 2A**). This strongly suggested the presence of coordinated activity in the same cell, however the correlation is lost when comparing HES1 and HES5 expression in different cells (**Supplementary Figure 2B**). Notably, dynamics with negative correlations were also observed albeit less frequently, corresponding to one HES having dominant expression while the other is very low (**Figure 3B, Supplementary Figure 2A**,B).

**Figure 3.**
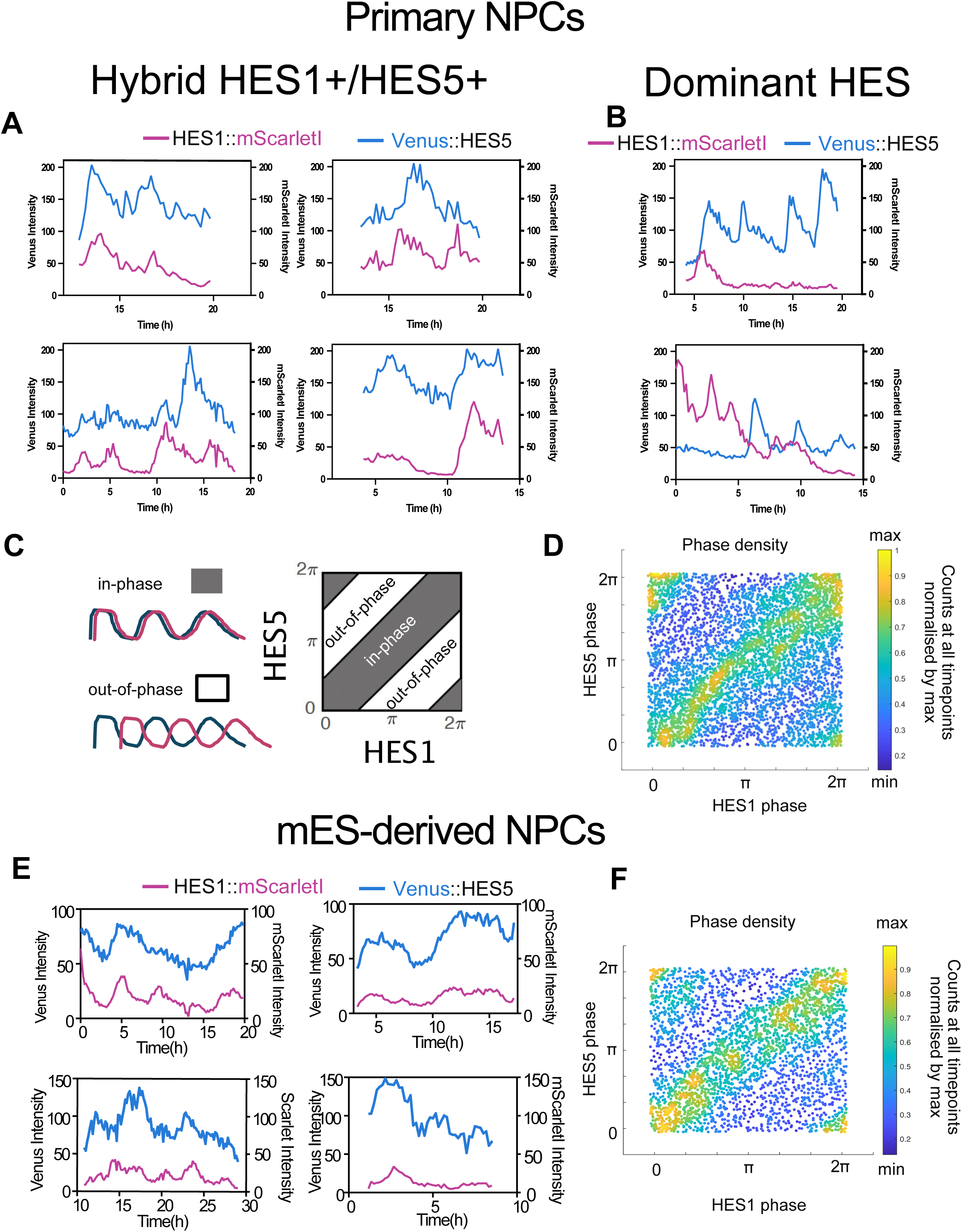
Exploration of synchrony in HES1 and HES5 oscillations in the same cells. **(A-B)** Representative examples of HES1::mScarlet-I (magenta) and Venus::HES5 (teal) raw intensity timeseries in the same cell observed in priNPC cultures in the same experiment grouped into co-expressing (or hybrid HES1+/HES5+ in **A**) and dominant HES in **B**; fluorescent units of Venus and Scarlet are not directly comparable, however the same fluorophore intensity can be compared across examples. **(C)** Diagram depicting areas of HES1 versus HES5 in-phase and out-of-phase regions of phase-phase density mapping in the same cell at the same time. **(D)** Phase-phase density mapping showing HES1 and HES5 phase values observed in the same cells at the same time in timeseries collected from the primary neural stem cells; markers are color-coded to indicate low and high probability density areas. **(E)** Representative examples raw timeseries of HES1-mScarlet-I and mVenus::HES5 observed in mES-derived NPCs. **(F)** Phase-phase density mapping showing HES1 and HES5 phase values observed in timeseries collected from mES-derived NPCs; markers are color-coded to indicate low and high probability density areas.

The predominance of positive correlation values in the same cell were consistent with the predominance of the hybrid HES1+/HES5+ cell state in fixed timepoint in-vitro observations (**Figure 1D,E**) and in addition, strongly suggested synchronisation of HES oscillations. To confirm synchrony, we analysed the phase relationship of HES1 and HES5 oscillations in the hybrid HES1+/HES5+ cell state, where phase refers to the position within the oscillation cycle at any given time (**Supplementary Figure 2C**). We used phase angle reconstruction using previously described techniques (Biga et al., 2021) and plotted the phase of HES1 versus the phase of HES5 in the same cell at the same time across all timeseries. In these phase-phase maps, synchronous in-phase oscillations generate data clustered along the first diagonal and at the opposite corners (**Figure 3C**). Phase-phase maps of HES1 and HES5 from priNPCs showed that despite the presence of stochasticity, the HES1 and HES5 oscillations are highly synchronous in priNPCs at single cell level (**Figure 3D**). In the mES-derived NPCs, HES1 and HES5 predominantly co-expressed and showed positively correlated dynamics with some examples of dominant HES5 (**Figure 3E**, last panel dominant HES5). Here as well, there was strong evidence of in-phase activity when HES1 and HES5 are co-expressed in the same cell (**Figure 3E,F**).

Taken together, the data indicates that HES1 and HES5 oscillate in-phase in the hybrid HES1+/HES5+ cell state thus providing a way to balance the expression of two TFs in a dynamic way over time. This prompted the need for understanding the mechanism leading to synchronisation.

### A coupled HES1-HES5 oscillation model explains synchronisation and suggests an elongation of the endogenous HES1 period through entrainment

Our in vitro data showed that the expression of HES1 and HES5 synchronise and adopt a similar 3-4h period in co-expressing cells. However, HES1 and HES5 have reported differences in kinetic parameters which can impact on periodicity. Specifically, while the mRNA half-life of HES1 and HES5 is very similar (between 20 to 30min), the protein half-life of HES1 is reported to be 22min which is considerably shorter than 80-90min reported for HES5 (Hirata et al., 2002, Bonev et al., 2012, Manning et al., 2019). To understand how protein stability differences may impact on the period duration, we used a previously developed model of HES1 protein repressing *Hes1* mRNA via a Hill functions with a set time delay of 29min and a Hill coefficient of 5, which produces oscillations of 2-3x in amplitude (that decay over time as the model is deterministic) (Lewis, 2003, Monk, 2003, Goodfellow et al., 2014). Using model parameters based on experimentally derived HES1 or HES5 mRNA and protein degradation rates (see Materials and methods), the HES model simulations indicated that the HES1 predicted period was approx. 2.5h whereas the HES5 predicted period was longer (3-4h range) (**Figure 4A; Supplementary Figure 3A**). However, in our cell culture data, the HES1 period was elongated to a mean of 3.3 hrs and similar to the HES5 period. Taken together these findings suggested that in NPC cultures, the HES1 oscillator is operating outside of its normal dynamic regime and has an elongated period that cannot be explained from the current reported values for protein and mRNA stability.

**Figure 4.**
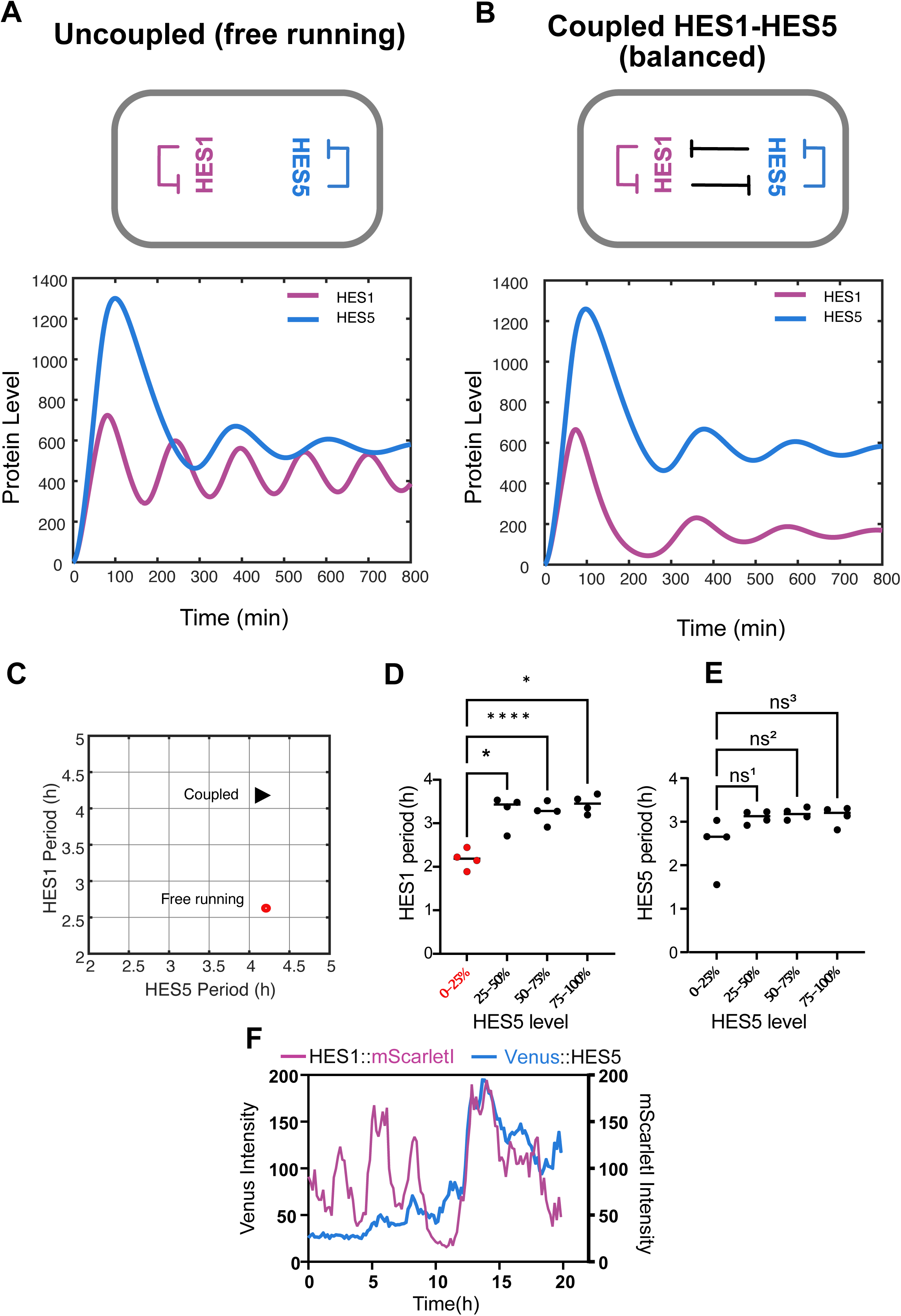
A mathematical model of coupled HES1-HES oscillations reveals entrainment of HES1 to the HES5 oscillatory period. **(A)** Simulation of free running (uncoupled) HES1 and HES5 dynamics at fixed values of period and mRNA degradation (Materials and methods); HES1 rates: 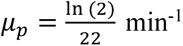, 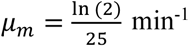; HES5 rates: 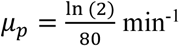, 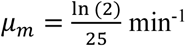; other model parameters: *α_m_* = *α_p_* = 1 min^-1^; *τ* = 29 min; *n* = 5;*P*_0_ = 390; **(B)** Simulation of coupled HES1 and HES5 dynamics at the same values of cross-repression (HES1 onto HES5; or HES5 onto HES1) and self-repression threshold values (HES1 onto HES1; HES5 onto HES5), referred to as balanced coupling *P*_015_ = *P*_051_ = *P*_01_ = *P*_05_ = 390 ; HES1 rates: 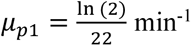, 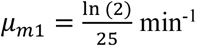; HES5 rates: 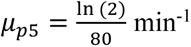, 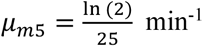;other model parameters: *α_m_*_1_ = *α_m_*_5_ = *α_p_*_1_ = *α_p_*_5_ = 1 min^-1^; *τ* = 29 min; *n* = 5. **(C)** Period duration (measured as average peak to peak interval) observed in free running versus coupled conditions. HES1 period elongates from approx. 2.6h to 4.1h whereas the HES5 period remains similar 4.2h versus 4.1h. **(D-E)** Median HES1 and HES5 period observed in primary neural progenitor cells with different level of HES5; binning of HES5 levels between 0 and 100% percentile values was performed and the median of HES5 period in those cells was computed per experiment; repeated measures ANOVA with Dunnet’s multiple comparison correction ns^1^=0.2989; ns^2^=0.1866; ns^3^=0.1169. **(F)** Representative example of priNPC showing a transition from single HES1 expressing to HES1+/HES5+ co-expressing over time; the HES1 period increases in the double expressing state.

We hypothesised that the HES1 period elongation is due to HES1-HES5 cross-repressive interactions causing entrainment. To test this hypothesis, we generated a coupled oscillators model that combines the effects of HES1-HES5 cross-repression in addition to self-repression (Materials and methods) and uses experimentally determined mRNA and protein degradation rates (Hirata et al., 2002, Bonev et al., 2012, Manning et al., 2019). In the coupled model repressive Hill functions are used to encode the cross-repression of one HES onto the other, referred to as H15 (HES1 protein repressing *Hes5* mRNA) and H51 (HES5 protein repressing *Hes1* mRNA), see Materials and methods. Both Hill functions introduce new parameters including time delays (set to 29min, same as auto-repression time delay), Hill coefficient (set to 5) as well as repression thresholds (P15 and P51 corresponding to H15 and H51 respectively) which are critically important in this system as they are inversely related to the strength of the cross-repression. The cross-repression threshold is defined as the amount of protein required to repress mRNA of the other gene by 50% (Monk, 2003). When the HES5 level is below the cross-repression threshold, HES1 is unaffected and remains subject to its own oscillatory dynamics. However, when HES5 levels reach above the repression threshold, cross-repression occurs. The same is true for HES1 cross-repression of HES5.

We compared dynamics observed in free running (uncoupled) HES1-HES5 conditions versus the coupled HES1-HES5 dynamics at equal values of P15/P51, a regime we refer to as balanced coupling (**Figure 4A vs B**). In the free-running conditions, the period of HES1 is shorter than HES5 as expected due to the relative differences in protein half-life. However, when the two are coupled they become entrained resulting in an elongated HES1 period similar to that of HES5 (**Figure 4C**). In other words, the slow oscillator HES5 cannot speed up and hence it wins the pace by forcing the fast oscillator, HES1, to slow down. We tested the existence of this phenomena in our priNPC data by comparing HES1 period in cells with different HES5 levels. Specifically, we binned the data into quartiles of average HES5 expression per cell, thus generating a total of 4 groupings (**Figure 4D,E**). In the lowest HES5 quartile group (0-25%, containing timeseries with low/no HES5) we observed that the period of HES1 is significantly shorter compared to other groups (1.6x reduced) (**Figure 4D**). This indicated that only 25% of maximum HES5 expression is sufficient to impact HES1 dynamics, consistent with the observation that in most cells (expressing above 25% HES5) the expression of both HES proteins adopt a similar period. As expected, the HES5 period is not significantly affected across the groupings despite becoming more variable at low HES5 where self-repression is likely impaired (**Figure 4E**). These results indicate that the presence of HES5 beyond a 25% level is sufficient to induce an elongation of HES1 period without perturbing HES5. Consistent with this, in some priNPC examples, we observed that the HES1 period can transition from short to elongated as HES5 becomes upregulated in the same cell (**Figure 4F**).

### Either HES can dominate at single cell level through a change in cross-repression threshold and/or level

As in other models of cross-repressive TFs, we also observed dynamic regimes where one HES can dominate over the other, which represent the ‘traditional’ bistable states where expression of one TF is maintained at a high level while the other is repressed over time (**Figure 5A,B** and **Supplementary Figure 3B**,C). These occurred at unbalanced coupling conditions where cross-repressive threshold values P15 and P51 respectively are lower than the thresholds of other interactions. In our simulations, strength is inversely proportional to the values of cross-repression threshold such that a lower threshold ensures that the dominant HES can rise above that value more easily and thus repress the other HES. When the strength of interaction from the dominant HES is higher compared to other interactions, this lack of balance results in either HES1 or HES5 switching the other off after a single pulse (**Figure 5A,B**). The effect of a change in cross-repression threshold is incremental as increased HES5 cross-repression leads to progressively less HES1 in the same cell through a severe dampening of HES1 peaks until only the initial one is observed (**Supplementary Figure 3B**). A similar effect can be obtained for a fixed cross-repression threshold value, in cases when the HES5 level is increased through higher mRNA and protein production rates leading to incremental suppression of HES1 peaks, albeit less effective in eliminating low amplitude pulsing compared to cross-repression (**Supplementary Figure 3C vs B**). We further explored how HES5 can dominate over HES1 through a combination of cross-repression and production rates. Indeed, the combined strategy is also effective in reducing maximum HES1, with the cross-repression threshold providing finetuning of HES5 onto HES1 repression even at low production rates, i.e. less than 1 (**Figure 5C**).

**Figure 5.**
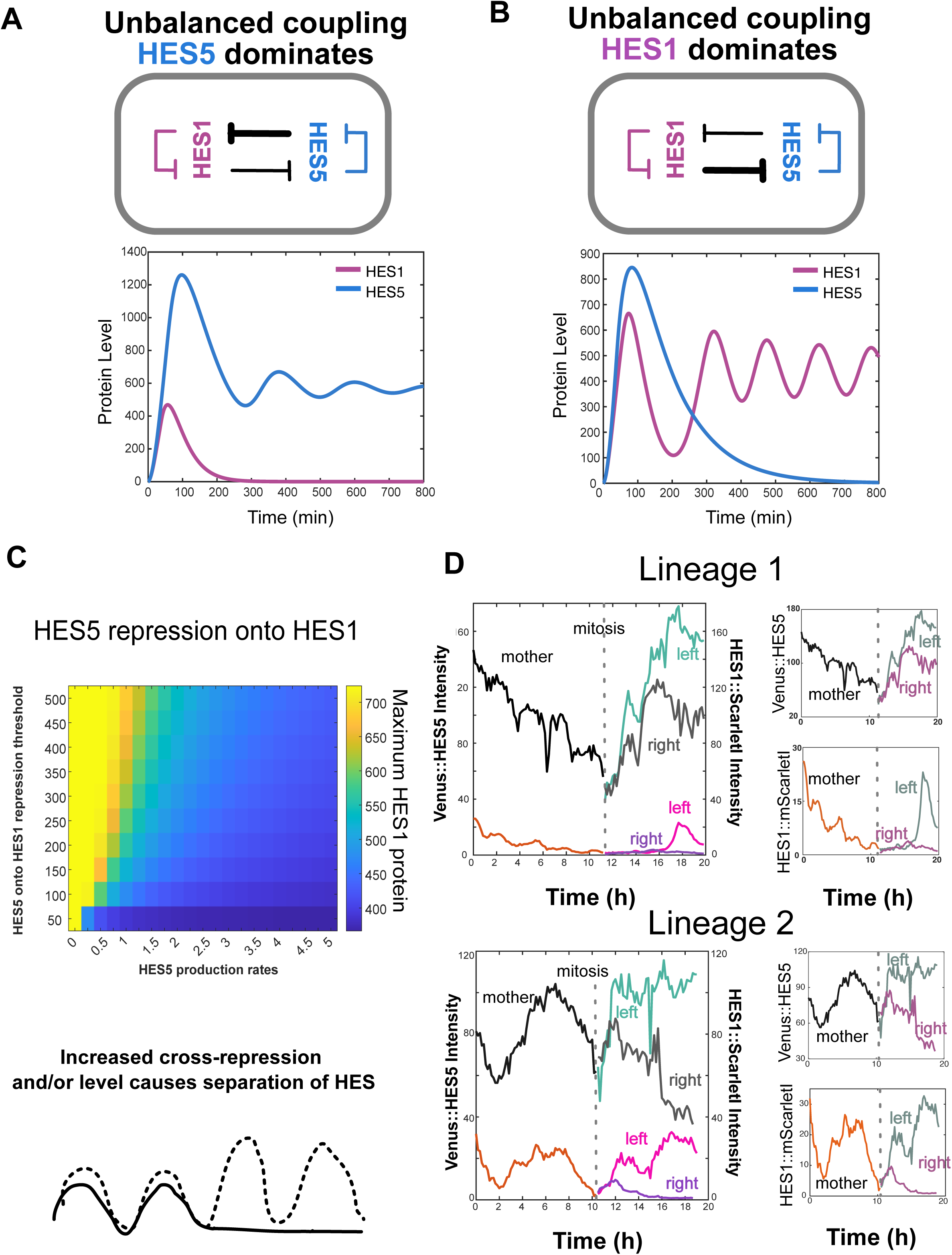
Mathematical modelling suggests unbalanced cross repression between HES1 and HES5 leads to a single dominant HES at single cell level. **(A-B)** Simulation of coupled HES1 and HES5 dynamics observed at cross-repression values that are unbalanced between HES1 and HES5 **(A)**, *P*_015_ = 390; *P*_051_ = 50 and **(B)**, *P*_015_ = 50; *P*_051_ = 390 respectively; these dynamic regimes allow one oscillator to supress the other after a single peak; following downregulation, HES revert to oscillations of different periods driven by self-repression. Model parameters: 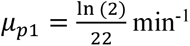, 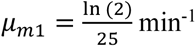; 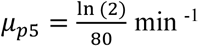, 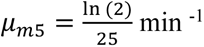, *α_m_*_1_ = *α_m_*_5_ = *α_p_*_1_ = *α_p_*_5_ = 1 min^-1^; *τ* = 29 min; *n* = 5. **(C)** Mapping of maximum HES1 protein level for a range of HES5 production rates (*α_m_*_5_ = *α_p_*_5_ ∈ [0,5]) and cross-repression threshold values (*P*_051_ ∈ [50,500]). Model parameters: *P*_015_ = *P*_01_ = *P*_05_ = 390, 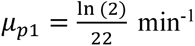, 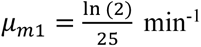; 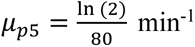, 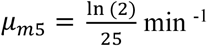, *α_m_*_1_ = *α_p_*_1_ = 1 min^-1^; *τ* = 29 min; *n* = 5. **(D)** Representative examples from mES-derived NPCs showing dynamics of HES1+/ HES5+ in mother and corresponding daughter cells (left and right); lineage 1&2: mother cell and left daughter co-express HES1 and HES5 whereas right daughter transitions into HES5+/HES1-.

Experimental observations from the mES-derived in-vitro cultures show that the transition from a co-oscillating hybrid state into a dominant HES state does occur in NPCs as they undergo neuronal differentiation (**Figure 5D**). Interestingly this seems to occur following divisions (**Figure 5D**) however, as expected, not all divisions have divergent HES patterns, with other examples showing co-expression can persists in both daughter cells (**Supplementary Figure 3D**). This shows that exit from the oscillatory HES1+/HES5+ hybrid state occurs at different times in progenitors in the same conditions.

The simulated mutually exclusive dynamic regimes, supported by experimental observations, suggest that HES1 and HES5 have the capacity to either co-express (in which case they co-oscillate) or resolve into one dominating over the other. Our model explorations show that exclusivity can occur through a change in repression threshold and/or increased abundance via production rates. In addition, experimental evidence suggests parameters can change in some of the progenitors following cell division.

### The HES1+/HES5+ hybrid state resolves during spinal cord development

To understand if transitions out of the hybrid state occur during development in vivo, we explored HES1 and HES5 expression domains along the mouse spinal cord dorsoventral axis. Previous reports have shown that at E10.5 HES5 encompasses separate ventral and dorsal expression domains, whereas HES1 is expressed dorsally (in areas where HES5 is low) as well as at the roof, floor plate and in p3 cells (situated above floor plate) (Sagner et al., 2018).

To identify if *Hes1* and *Hes5* may be co-expressed earlier in development and interact at single cell level, we used existing single cell transcriptomics data from mouse embryonic spinal cord (Delile et al., 2019). Mapping of level and percent expressing progenitors along the dorso-ventral axis between E9.5 and E11.5, shows that both *Hes* genes are widely found in spinal cord progenitors with a tendency for *Hes5* to become more abundant over development and *Hes1* to become more restricted to areas where *Hes5* is low over time (**Figure 6A**). The biggest changes are observed between E9.5 and E10.5 with little re-arrangement of *Hes* expressing domains at E11.5 (**Figure 6A**). This suggests it is possible for the two HES proteins to co-express at early stages (E9.5) in areas where neither HES1 or HES5 dominates and for co-expression to reduce at later stages as the domains become established (E10.5 onward).

**Figure 6.**
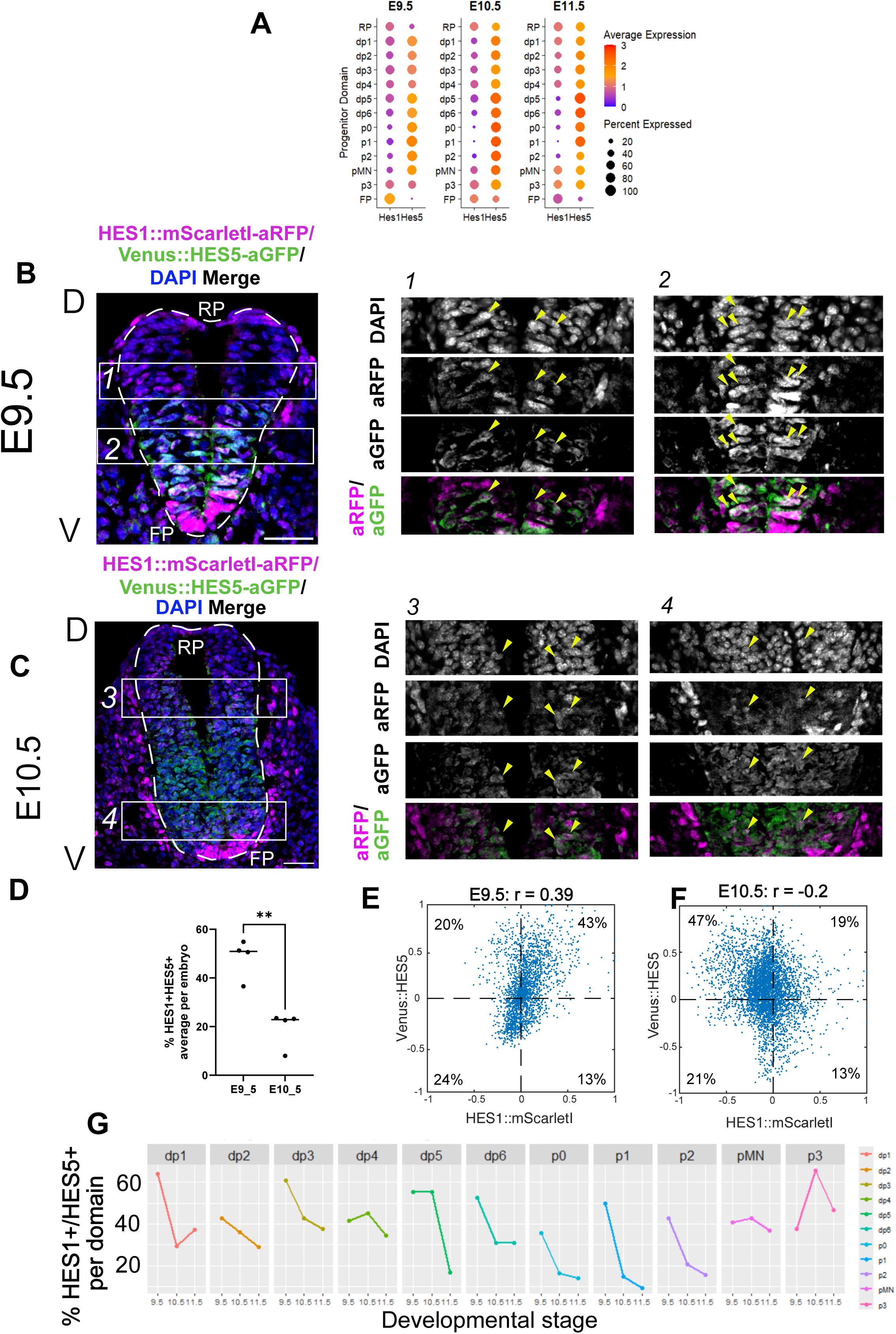
Spinal cord tissue expression of HES1 and HES5 at different stages in development. **(A)** Quantification of *Hes1* and *Hes5* expression in progenitors in the dorso-ventral axis of mouse embryonic spinal cord between E9.5 to E11.5 based on **t**ranscriptomic data from (Delile et al., 2019). **(B-C)** Cryosections of mouse embryonic spinal cord at E9.5 **(B)** and E10.5 **(C)** in embryos containing endogenous HES1::mScarlet-I and Venus::HES5 imaged at single cell resolution; color channels represent immunofluorescence of nuclear DAPI, HES1::mScarlet-I detected with anti-RFP and Venus::HES5 detected with anti-GFP in the same slice (Materials and methods); HES1 is highly expressed at the floor plate (FP) and roof plate (RP) where no HES5 is present; high-magnification inset areas (*1-4*) depict double expressing nuclei (arrowheads). **(D)** Comparison of E9.5 versus E10.5 percentage of nuclei co-expressing HES1::mScarlet-I (from anti-RFP) and Venus::HES5 (from anti-GFP); nuclei found within the basal boundary were considered excluding cells in RP and FP (identified as HES1+); markers indicate mean per experiment from multiple cryosections; unpaired t-test, 2-tailed with p<0.01. **(E-F)** Scatter plot representation of single cell data collected from E9.5 **(E)** and E10.5 **(F)** cryosections in nuclei within the basal boundary excluding RP and FP; intensity values were background subtracted and normalised to maximum per section (Materials and methods); Pearsons correlation coefficient (r); percentages in each quarter correspond to fractions of single, double and negative expressing nuclei observed overall. **(G)** Co-expression analysis using transcriptomics data in **(A)** in different progenitor types in the dorso-ventral axis.

To confirm these findings, we used cryosections of dual HES1::Scarlet-I/Venus::HES5 spinal cords at E9.5 and E10.5 (**Figure 6B, C**). We used antibodies against mScarlet-I (anti-RFP) and Venus (anti-GFP) to increase the signal to noise ratio and detect nuclear expression of the HES proteins more accurately (Materials and methods). We observed that HES1 expression is highest at the floor plate and also present at the roof plate where no HES5 is observed (**Figure 6B,C** left-most panel). Outside of the floor plate and roof plate regions, we observed that at E9.5, HES5 showed a prominent ventral expression domain, with HES1 expression overlapping both the dorsal and ventral sides of the domain, indicating co-expression of HES1 and HES5 in the dorsoventral axis **(Figure 6B**). At E10.5, the HES1 dorsal and HES5 ventral expression domains became larger comprising more nuclei and areas of co-expression of HES1 and HES5 became further restricted to the edges of the HES5 ventral domain (**Figure 6C**). Using a stringent intensity threshold and DAPI to identify nuclei of progenitors within the neural tube, we quantified the percentage of cells expressing HES1 and HES5 across the dorsoventral axis but excluding the floor and roof plate regions (**Supplementary Figure 4A**,B and Materials and Methods). We consistently observed that the spinal cord of early E9.5 embryos contained approx. 50% double positive nuclei however this is reduced to only about 23% double positive nuclei at E10.5 (**Figure 6D**).

Consistent with this, we observed that while nuclear intensities at E9.5 show a weak positive correlation in HES1 versus HES5, at E10.5 this is lost primarily through an increase in the HES5 only fraction at the expense of double positives **(Figure 6E,F**). These observations suggested the ventral HES5 only expression domain becomes more established during development, which in turn leads to a reduction of areas where HES1 and HES5 can overlap. Our analysis of existing transcriptomics data (Delile et al., 2019) supports the observation that co-expressing *Hes1+*/*Hes5+* fractions become reduced over developmental time particularly in ventral spinal cord, however co-expression persists for longer in dp4-dp5 and p3 which are progenitor types observed either side of the ventral *Hes5* domain (**Figure 6G**).

Taken together, our in-vitro, tissue and theoretical observations suggest that cross-repressive TFs HES1 and HES5 are co-expressed at single cell level in early spinal cord development, forming a hybrid state where they oscillate in-phase, with HES5 entraining HES1 to a slower period. However, at later stages, through bistability one HES dominates over the other through cross-repressive interactions and most progenitors adopt one or the other HES. In our model, this effect can be explained through a reduction in the thresholds of cross-repression and/or increase in the level of one HES, see model in Figure 7.

**Figure 7.**
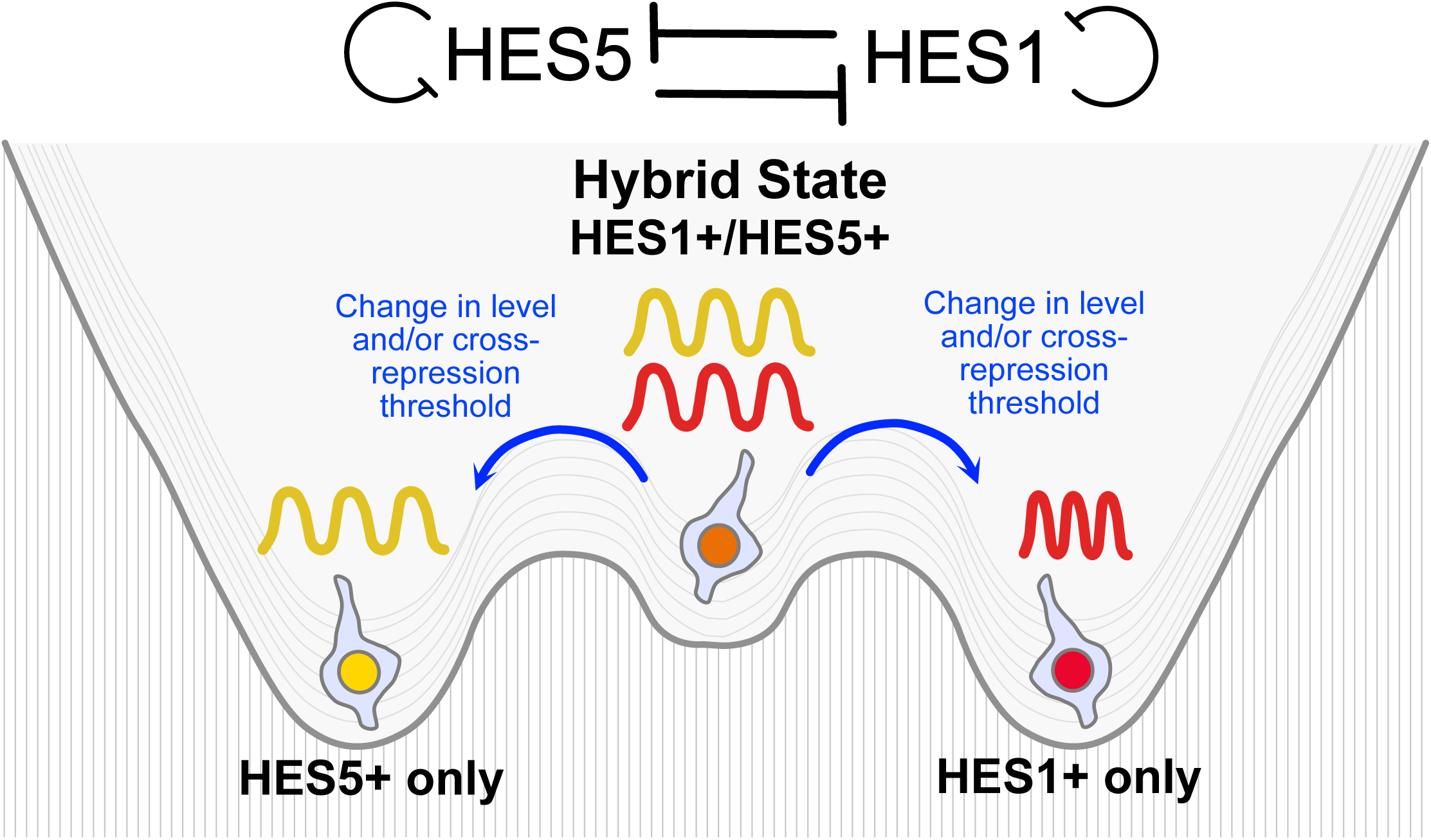
Graphical abstract.

## Discussion

Using in-vitro data, we show that HES1 and HES5 can co-oscillate in NPCs representing a hybrid state from which cells can transition into a single dominant HES. This is supported by in-vivo data indicating that a transition from the hybrid to a dominant HES state occurs in spinal cord over development. We surmise that the hybrid state represents a primitive stage because in vivo more cells co-expressed HES1 and HES5 at stage E9.5 than at stage E10.5. Analysis of protein (this study) complemented by existing transcriptomics data (Kicheva et al., 2014, Sagner et al., 2018, Delile et al., 2019) suggests that different spinal cord progenitor groups escape the hybrid state at different times, with dorsal progenitors appearing delayed compared to most ventral progenitor types suggesting that a spatio-temporal mechanism to control this process is required, and could contribute to the low rate of differentiation at early time points in the dorsal regions in the spinal cord in (Kicheva et al., 2014). Using computation, we demonstrate that both hybrid and dominant states can be explained by cross-repression between HES1 and 5. The resolution of cross-repressive and oscillatory TFs takes place in the model when cross-repression in either direction becomes stronger than the other, and stronger than autorepression, or when production is increased, or indeed a combination of the two strategies. Resolving gene expression through a hybrid, double-positive oscillatory state, rather than a bistable switch offers several advantages. One key benefit is the ability to extend the decision-making window. In some progenitor cells, this hybrid oscillatory state may delay the final resolution of the switch, allowing expression domains to be gradually established and refined over time, in sync with the timeline of neurogenesis. Additionally, this mechanism provides a form of functional compensation; under certain parameter conditions, the non-dominant HES can continue to oscillate at low amplitude, remaining poised for activation. If repression is lifted, for example, through deletion of the dominant factor, this latent activity could allow rapid upregulation across the spinal cord, supporting robustness in patterning (Hatakeyama et al., 2004).

Additionally, comparing the co-expression with that of single oscillatory HES1 or HES5 allowed us to gain some important insights of how two HES oscillators interact within single cells. We were able to establish that the free running period of HES1 is shorter than the free running period of HES5, as indeed is predicted by their differential protein stability. However, when co-expressed, in-phase oscillations of HES1 and HES5 with a common longer period are observed. Our computational modelling suggests an entrainment event whereby the slow oscillator, HES5, forces the fast oscillator, HES1, to slow down and as a result they acquire common periodicity and phase. This extends our understanding of frequency deformations beyond systems of identical oscillators (Rohm, 2018) and highlights the versatility of coupled oscillators, in this case enabling HES1 to oscillate outside of its normal regime, a behaviour suggesting nonlinearity close to an infinite period bifurcation such as observed in the somitogenesis clock (Sanchez et al., 2022).

Inter-cellular coupled oscillator systems can drive embryonic patterning, for example in somitogenesis Notch signalling synchronises HES7 oscillations between cells to ensure whole somites form in a rhythmic and ordered manner along the anterior-posterior axis (Niwa et al., 2007). More recently we showed that in the embryonic spinal cord, intercellular coupling of HES5 results in local synchronisation of oscillations posited to control the rate and spacing of differentiation events (Biga et al., 2021, Hawley et al., 2022, Hawley et al., 2025). Here, we showed that different oscillators can also be coupled within the same cell leading to their intra-cellular synchronisation. This prompts the question of what is the function or advantage of this intra-cellular coupling? Controlling the phase of oscillators is a powerful way to control their biological outcome. For example, in somitogenesis, the relative timing between Notch and Wnt oscillators (i.e phase shift) changes along the anterior-posterior axis of the pre-somitic mesoderm and that the phase-shift controls somite differentiation (Sonnen et al., 2018). In a different context, the phase divergence (or temporal redundancy) of TF paralogs with overlapping targets can be modulated from out-of-phase to in-phase to upregulate the level of target genes during yeast stress response (Wu et al., 2021). HES proteins are transcriptional repressors, and we have previously postulated that the time that a cell spends expressing a low level of HES (i.e. a trough in the oscillations), affects the probability of downstream genes being activated and neuronal differentiation initiated (Biga et al., 2021, Hawley et al., 2022). Single knock-out experiments (Hatakeyama et al., 2004) indicate that HES1 and HES5 can functionally compensate for each other suggesting that they have at least some common downstream targets. A random phase or an out-of-phase oscillation of HES1 and HES5 would lead to overall noisy high HES levels which in turn could supress pro-neural targets and thus block differentiation, see (Soto et al., 2020). In contrast, in-phase oscillations may promote robustness of Notch signalling response (via Delta-like 1) and precisely control the level of pro-neural targets without supressing them. Synchrony of paired zebrafish ortholog genes *her1* and *her7* promote robust pattern formation during somitogenesis (Zinani et al., 2021). Furthermore, in the zebrafish forebrain, we have shown that inter-cellular coupling of Her6 confers robustness against dynamic perturbations to maintain a normal phenotype (Doostdar et al., 2024), so by analogy it is also possible for intra-cellular coupling to increase robustness.

In conclusion, oscillatory expression enables a HES1 and 5 hybrid state which resolves into a single HES **(Figure 7)** leading to spatial segregation of HES1 and HES5 in spinal cord. In addition, coupling between these two oscillators and the resulting synchrony when they are expressed in the same cells, may preserve and even enhance the decoding functionality of HES oscillators.

## Materials and Methods

The list of reagents and antibodies is included in **Table 1**.

**Table 1.** List of reagents.

| Reagent or Resource | Source | Catalogue No. |
| --- | --- | --- |
| **Antibodies** |  |  |
| Mouse monoclonal anti-PAX3/PAX7 (1:200) | R & D Biotech | MAB2457 |
| Rabbit polyclonal anti-SOX2 (1:200) | Abcam | ab97959 |
| Anti-RFP (1:500) | Proteintech | 5F8 |
| Anti-GFP (1:200) | Abcam | ab13970 |
| Anti-GFP (1:100) | Thermo Fisher | A-11122 |
| Anti-GFP (1:500) | Roche | 11814460001 |
| Donkey anti-rabbit Alexa Fluor 488 (1:500) | Thermo Fisher Scientific | A21206 |
| Donkey anti-mouse Alexa Fluor 647 (1:500) | Thermo Fisher Scientific | A31571 |
| Donkey- anti chicken Alexa Fluor 488 (1:500) | Jackson ImmunoResearch | 703-545-155 |
| Donkey anti-Rat DyLight 550 (1:800) | Thermo Fisher Scientific | SA5-10027 |
| Goat anti-mouse Alexa Fluor 647 (1:500) | Thermo Fisher Scientific | A-21235 |
| Goat anti-rabbit Alexa Fluor 405 (1:500) | Thermo Fisher Scientific | 31556 |
| **Cell Culture and IF Reagents** |  |  |
| DMEM/F12 | Sigma | D6421 |
| Neurobasal | Gibco | 21103-049 |
| ESGRO 2i LIF | Sigma | SF016-100 |
| N2 supplement | Gibco | 17502048 |
| B27 supplement | Gibco | 17504-044 |
| GlutaMAX | Gibco | 35050-038 |
| D-Glucose solution | Sigma | G8644 |
| Bovine Albumin (BSA) Fraction V 7.5% | Gibco | 15260-037 |
| MEM Non-essential Amino Acid Solution | Sigma | M7145 |
| Phosphate buffer saline (PBS) | Sigma | D8537 |
| Basic fibroblast growth factor (bFGF) | PeproTech | 100-18B |
| Mouse EGF recombinant protein | PeproTech | 315-09-500UG |
| CHIR-99021 | Sigma | SML1046-5MG |
| LIF | Bio-Techne | 8878-LF-025 |
| PD0325901 | Sigma | PZ0162-5MG |
| Retinoic acid (RA) | Sigma | R2625 |
| Accutase | Sigma | A6964 |
| Laminin | Sigma | L2020 |
| Laminin 511 | Biolamina | LN511-0202 |
| Matrigel | Corning | 356234 |
| Donkey serum | Sigma | D9663 |
| Fibronectin | Millipore | FC010 |
| KSOM | Millipore | MR-101-D |
| Acid Tyrode's solution | Millipore | MR-004-D |
| Paraformaldehyde (PFA) | Thermo Fisher Scientific | 28908 |
| Triton X | Sigma | T9284-500ML |
| 2-mercaptoethanol | Invitrogen | 31350-010 |
| Rabbit anti-mouse antiserum | Sigma | M5774 |
| Guinea pig serum | Sigma | C5405 |
| Hoechst 33342 Solution | Thermo Fisher Scientific | 62249 |
| **Other** |  |  |
| CELLview 35-mm cell culture dish | Greiner BIO-ONE | 627860 |
| CELLview 35-mm cell culture dish (4 compartments) | Greiner BIO-ONE | 627870 |
| Superfrost Plus Adhesion microscope slides | Eprendia | J1800AMNZ |
| Gold Antifade | Thermo Fisher | P36930 |

### Mouse lines

Animal experiments were performed within the conditions of the Animal (Scientific Procedures) Act 1986. *Venus::Hes5* knock-in mice (ICR.Cg-Hes5<tm1(venus)Imayo>) (Imayoshi et al., 2013) were obtained from Riken Biological Resource Centre, Japan. *Hes1::mScarlet-I* knock-in mice were generated in (Marinopoulou et al., 2021). We used the EASI-CRISPR strategy (Quadros et al., 2017) to generate a C-terminally tagged *Hes1::mVenus* mouse line using the same protocol (Marinopoulou et al., 2021, Bennett et al., 2021). To generate the long single strand DNA donor repair template, a homology flanked flexible linker-3xFLAG-mVenus DNA sequence was cloned and used as a template in an initial PCR reaction with primers Hes1_lssDNA_F catgctcccggccgcCATGGGAATTCGGTACcaacagtgggacctcggt and Hes1_lssDNA_R caagttcgtttttagtgtccgtcagaagagagaggtgggctagggactttacgggtagcagtggcctgaggctctcacttgtacagctcgtccatgcc. Potential founder mice were screened by PCR, using primers that flank the homology arms (Geno F ttgcctttctcatccccaac, Geno R gcagtgcatggtcagtcac), used in combination with internal mVenus primers (mVenus F CACATGAAGCAGCACGACTT and mVenus R TCCTTGAAGTCGATGCCCTT). Germline transmission was confirmed through PCR and sequencing and a colony was established. For timed matings, E0.5 was considered as midday on the day a plug was detected.

### Establishment and maintenance of primary neural progenitor cultures

Primary neural progenitor cells were isolated from dissected spinal cords of E10.5 embryos obtained from crossing *Venus::Hes5* and *Hes1::mScarlet-I* knock-in mice or dissecting LGEs of E13.5 embryos from crossing the *Hes1::mVenus* with *Hes1::mScarlet-I* knock-in mice. All lines were genotyped to confirm presence of knock-in fusion alleles, mScarlet-I (Marinopoulou et al., 2021) and Venus/mVenus genotyping primers (Venus R CTACTTGTACAGCTCGTCCATGCC and Venus F GTGTCTAAGGGCGAAGAGCTG; mVenus as listed above), to identify cells expressing both fluorophores. Primary lines were maintained using protocols from (Pollard, 2013). Dissociated cells were cultured on Laminin coated plates in DMEM/F-12 media containing 4.5 mg/ml glucose, 1× MEM non-essential amino acids, 120 μg/ml Bovine Album Fraction V, 55 uM 2-mercaptoethanol, 1× GlutaMAX, 0.5× B27, 0.5× N2 supplemented with 10ng/ml EGF and 10ng/ml bFGF. Cell lines were passaged routinely every 2-3 days by dissociation with Accutase and re-seeded at split ratios between 1:2 to 1:4. For imaging experiments cells were seeded at 12,000 to 23,000 cells/cm^2^ in Cell view 35mm glass-bottomed dish and allowed to proliferate for 2-3 days.

### Establishment and maintenance of Venus::HES5/HES1::mScarlet-I mouse ES cell-lines

All incubation steps were carried out at 37°C, 5% CO2. ES cell derivations were performed as described in (Nichols et al., 2009) with small alterations: Embryos were flushed from oviducts at 4-8 cell stage (E2.5) and placed into KSOM plus 2i (1µM PDO325901 and 3µM CHIR99021) with PBS into the outer well to avoid evaporation, and incubated for 2 days. If the embryos had not hatched, the zona pellucida was removed using Acid Tyrode’s solution, and the trophectoderm was removed by incubating for 1 hour the embryos in KSOM + 20% Rabbit anti-mouse antiserum, rinsed 3x in pre-equilibrated N2B27 media (comprised of 1:1 mix of Neurobasal and DMEM/F12 plus 1x Glutamax, 0.5X N2, 0.5X, 0.5% BSA and 0.1mM β-mercaptoethanol) and then incubation for 30mins in KSOM + 20% Guinea pig serum. The trophectoderm lysate was then manually removed using a finely drawn pipette. The isolated ICM cells were placed into separate wells of a 96 well plate (pre-coated with 20µg/ml Laminin 511 + 1mM Fibronectin in N2B27+2i + 10ng/ml LIF, and incubated for approximately 7 days, until there is a large outgrowth. The outgrowths were passaged using Accutase and gradually expanded until the lines were established, and at which point were maintained on 0.2% gelatin coated plates in N2B27+ 2i + LIF (Ying et al., 2008) or in ESGRO+2i LIF.

### Differentiation of mouse ES cells to dorsal neural progenitors

We used a previously described protocol that enriches for dorsal interneurons 4 to 6 (Gupta et al., 2022). The differentiation media consisted of 1:1 DMEM/F-12 and Neurobasal supplemented with 0.5xN2, 1xB27, 2mM Glutamax, 0.1mM 2-mercaptoethanol and 1%BSA. Between 50,000 to 100,000 mES cells per cm2 were seeded on Matrigel coated culture Cell view 35mm glass-bottomed dish with 4 partitions in differentiation media containing 10ng/ml bFGF (day1) followed by 10ng/ml bFGF and 3µM CHIR99021(day2) to generate neuromesodermal precursors (Gouti et al 2014). Neural induction was carried out with 100nM RA (days 3-4). From day 5 onwards cells were maintained in differentiation media (no RA) and Venus-HES5 expression was observed after 1-2 days.

### Timelapse imaging and single cell tracking

Movies were acquired using Zeiss LSM880 microscope and GaAsP detectors using a Fluar ×40 objective. HES1-mScarlet and HES5-Venus were detected using a 561nm and either 488nm or 514nm laser (Venus). was de Z-sections covering the depth of the cells were acquired every 10min. Primary neural stem cells were tracked using maximum intensity projections images created with Fiji. Single cell tracks were produced in Imaris using the ‘Spots’ and ‘Track over time’ function using the Brownian motion algorithm. All tracks were manually curated to ensure accurate single cell tracking, and generated mean intensity over time, representing protein concentration over time in single cells. The 2D spot size was set to 2.5um diameter in X and Y dimensions. Tracking of mES-derived neural progenitors was performed in 3D images to account for the fact that individual cells can overlap in the z-plane. The 3D spot size used was 3um diameter in X,Y and Z dimensions.

### Immunofluorescence of cryosections

Mouse embryo trunks from E9.5 and E10.5 embryos were fixed in 4% PFA for 1 h at 4°C, followed by three quick washes with 1xPBS and one longer wash for 1 h at 4°C. Embryos were equilibrated overnight in 30% sucrose (MERCK) at 4°C before mounting in Tissue-Tek OCT (Sakura) in cryomoulds and freezing at −80°C. 12 µm sections were cut on Leica CM3050S cryostat and collected using Superfrost Plus adhesion microscope slides. For immunofluorescence, sections were washed twice with PBS and stained using the same protocol as for cultured cells. Cells were fixed using 4% PFA for 15 minutes followed by a PBS wash. Permeabilisation and blocking was carried out using PBS + 0.1% Triton X + 10% donkey serum for 1 hour at room temperature. Primary antibodies (see **Table 1**) were diluted in PBS + 0.1% Triton X + 1% donkey serum and incubated on the samples overnight at 4°C. Three 10 minutes washes in PBS was then performed and then secondary antibodies (see **Table 1**) added, diluted in PBS + 0.1% Triton X + 1% donkey serum, incubated at room temperature for 4 hours (for tissue sections), kept in the dark or left at 4C overnight (for cultured cells). Nuclei were stained with DAPI or Hoechst 33342. Three more PBS washes were then performed, and the cells left in PBS for imaging, or tissue sections were mounted with Gold Antifade Mountant. For tissue sections, anti-RFP antibody was used to detect Hes1::mScarlet-I endogenous knock-in, and three GFP antibodies were used to detect Venus::HES5 all with similar results.

### Quantification of positive cells

Endogenous levels of HES1-mScarlet and HES5-Venus were quantified as mean intensity per nucleus using the Imaris ‘Spots’ function applied to the DAPI/Hoechst nuclear marker channel. Negative mean intensity data was generated from cell cultures not containing Venus or mScarlet-I-I which were processed in the same way. Mean intensities above the 95^th^ percentile of negative control values were considered positive for HES1::mScarlet-I and Venus::HES5. Immunofluorescence data from cell cultures and tissue was quantified in the same way, ie mean intensity per nuclear spot (detected from DAPI/Hoechst). In cultures, the 95^th^ percentile of values observed secondary only control samples was used as a threshold for positive cells. In tissue, positive nuclei values were considered above the 95^th^ percentile of mean intensities of GFP-/RFP- nuclei located outside the neural tube per slice. The criterion used for tissue is more stringent compared to a secondary only control (**Supplementary Fig4 A,B**).

### Detection of oscillations by a Gaussian Process method

We used the statistical approach developed by (Phillips et al., 2017, Manning et al., 2019) to analyse periodicity in single cell timeseries. Briefly, data was de-trended using a 7.5 lengthscale parameter to remove long-term behaviour, such as down-regulation and to recover the oscillatory signal with zero mean. We used maximum-likelihood estimation to fit the de-trended data timeseries with two competing models: a fluctuating aperiodic one (null model) and an oscillatory one (alternative model). We used the log-likelihood ratio (LLR) statistic to compare the likelihood of data being oscillatory or non-oscillatory and determined the oscillators based on a false discovery rate of 5% independently per experiment.

### Peak to trough fold change

We used the routines described in (Manning et al., 2019) to identify peaks and troughs in the timeseries. Peaks were paired with subsequent troughs and we computed peak to trough fold change as the ratio of intensity at the peak divided by intensity at the trough per track.

### Phase-phase mapping

The Hilbert transform was used to reconstruct the phase angle in the HES1 and HES5 timeseries. Phase-phase mappings were generated by plotting the phase of HES1 against the phase of HES5 in the same cell at any time. We used dscatter.m (Eilers and Goeman, 2004) to color code the density of datapoints.

### Mathematical modelling

We used the deterministic model of HES mRNA-protein interactions described in (Lewis, 2003, Monk, 2003) to model the variation of protein (P) and mRNA (m) over time dependent on production, degradation rates and delayed self-repression:

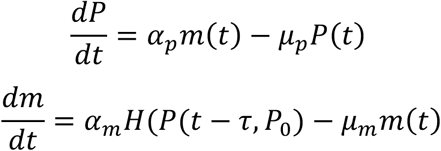

where *α_p_*, *α_m_* translation, transcription (in absence of protein repression) rates and *μ_p_*, *μ_m_* represent degradation rates for protein and mRNA respectively. The interaction between protein and mRNA occurs through a repressive Hill function:

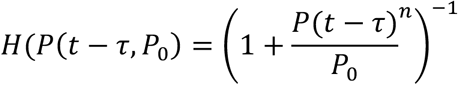

where *τ* represents time delay, *P*_0_ represents repression threshold and *n* is the Hill coefficient.

We selected values for the parameters that generate persistent oscillations of HES, consistent with previous reports (Goodfellow et al., 2014, Lewis, 2003, Monk, 2003). Values of *α_p_*, *α_m_* were set to 1 min^-1^(Goodfellow et al., 2014).The auto-repressive time delay *τ* was set to 29 min (Lewis, 2003); repression threshold *P*_#_ was set to 390, a value identified to oscillate in (Goodfellow et al., 2014); the Hill coefficient n = 5, a value that yields persistent 2x to 3x peak to trough amplitude of oscillations with minor decay (Monk, 2003) similar to what we have observed in the data.

We estimated the period (measured as average peak-to-peak interval) that emerges at different stability of mRNA and protein by taking into account that degradation rates are related to half-life, 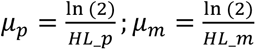 where *HL_p_*, *HL_m_* represent protein and mRNA half-life respectively. In Supplementary Figure 3 we can observe that the predicted period for HES1 is shorter than for HES5 primarily due to large differences in reported protein half-life (22min for HES1 (Hirata et al., 2002); 80-90min for HES5 (Manning et al., 2019)) and to some extent also due to small differences in mRNA half-life (25min for Hes1 (Bonev et al., 2012); 30min for Hes5 (Manning et al., 2019)).

We then generated a coupled model of HES1 and HES5 including delayed auto-repression as well as delayed cross-repression:

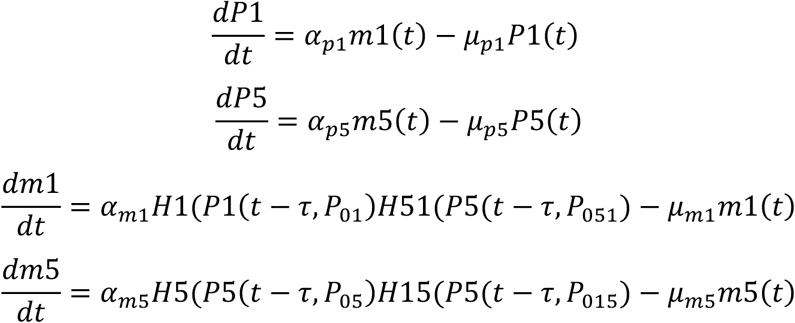

where *P*1, *m*1 and *P*5, *m*5 correspond to HES1 and HES5 protein, mRNA respectively. In addition to parameters found in the single HES model, the coupled version contains *H*1, *H*5 auto-repressive Hill functions for HES1 and HES5 as well as cross-repressive Hill functions *H*15, representing repression from the HES1 protein onto Hes5 mRNA and *H*51, representing repression from the HES5 protein onto Hes1 mRNA. Each Hill function has characteristic parameters: time delays, *τ* = 29 min; Hill coefficient *n* = 5 and auto-repressive thresholds *P*_01_, *P*_05_ = 390 and cross-repressive thresholds *P*_015,_*P*_051_ that were varied from 390 to 50 in different simulations (Figures 4-5). Repression thresholds are inversely proportional to repression strength such that a low value indicates higher repression compared to a high value.

Other parameters were set in a similar way to the single HES model. Translation and transcription rates in absence of protein repression) were set to *α_p_*_1_ = *α_m_*_1_ = *α_p_*_5_ = *α_m_*_5_ = 1 min^-1^ and identical to the single HES model. The protein degradation rates were set to 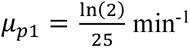 and 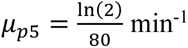 corresponding to 25min and 80min half-life values for HES1 and HES5 protein respectively. As the stability of mRNA is generally low for both HES (Bonev et al., 2012, Manning et al., 2019), the mRNA degradation rates were set to 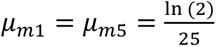 corresponding to 25min half-life for either HES resulting in HES5 periods similar to what was observed in the data.

### Statistical analysis

Statistical analysis was performed using Prism 10.3.0 with details in Table 2. Detection of periodicity, peak to trough fold change and phase angle was carried out using custom Matlab R2020a routines described in (Phillips et al., 2017, Manning et al., 2019, Marinopoulou et al., 2021). Mathematical modelling will be made available in Github.

### Bioinformatic analysis

sc-RNAseq data from (Delile et al., 2019) was analysed in R using Seurat v5 (Hao et al., 2024). Cells were excluded if they had more than 6% UMI counts associated with mitochondrial genes and expressed less than 800 genes. Only progenitor cells (annotated in Delile et al) and E9.5, E10.5 and E11.5 time points were used for the analysis. Co-expression of *Hes1* and *Hes5* was assigned to cells if they had at least 1 count for both genes.

## Acknowledgements

The authors thank the members of the Manning, Papalopulu and Francois labs as well as Andreas Sagner, Raman Das, Shane Herbert and Holly Lovegrove for useful discussion. We also thank the following for technical assistance, Robert Lea in Manning Lab, David Spiller, University of Manchester Bioimaging, Biological Services and the Genomics Unit.

## Author Contributions

Conceptualization: VB, CM, NP, PF, AM. Methodology: VB, AM, EM, AA, CM. Software: VB, AM. Validation: VB, AM, EM, AA. Formal analysis: VB, AM, AK, YM. Investigation: VB, AM, AK, YM. Resources: VB, AM, NP, EM, CM. Data curation: VB, AM, CM. Writing-original draft: VB, CM, NP, AM. Writing-review and editing: VB, CM, NP, AM. Visualisation: VB, AM, AK, YM. Supervision: VB, CM, NP, AM. Project administration: VB, NP, CM. Funding acquisition: VB, NP, CM.

