## Supplementary Figures and Legends for "Oscillatory Co-expression of HES1 and HES5 Enables a Hybrid State in a Bistable Transcription Factor Regulatory Motif"

**Supplementary Figures- pages 2 to 5**

**Supplementary Figure Legends- pages 6 to 7**

### Supplementary Figure 1

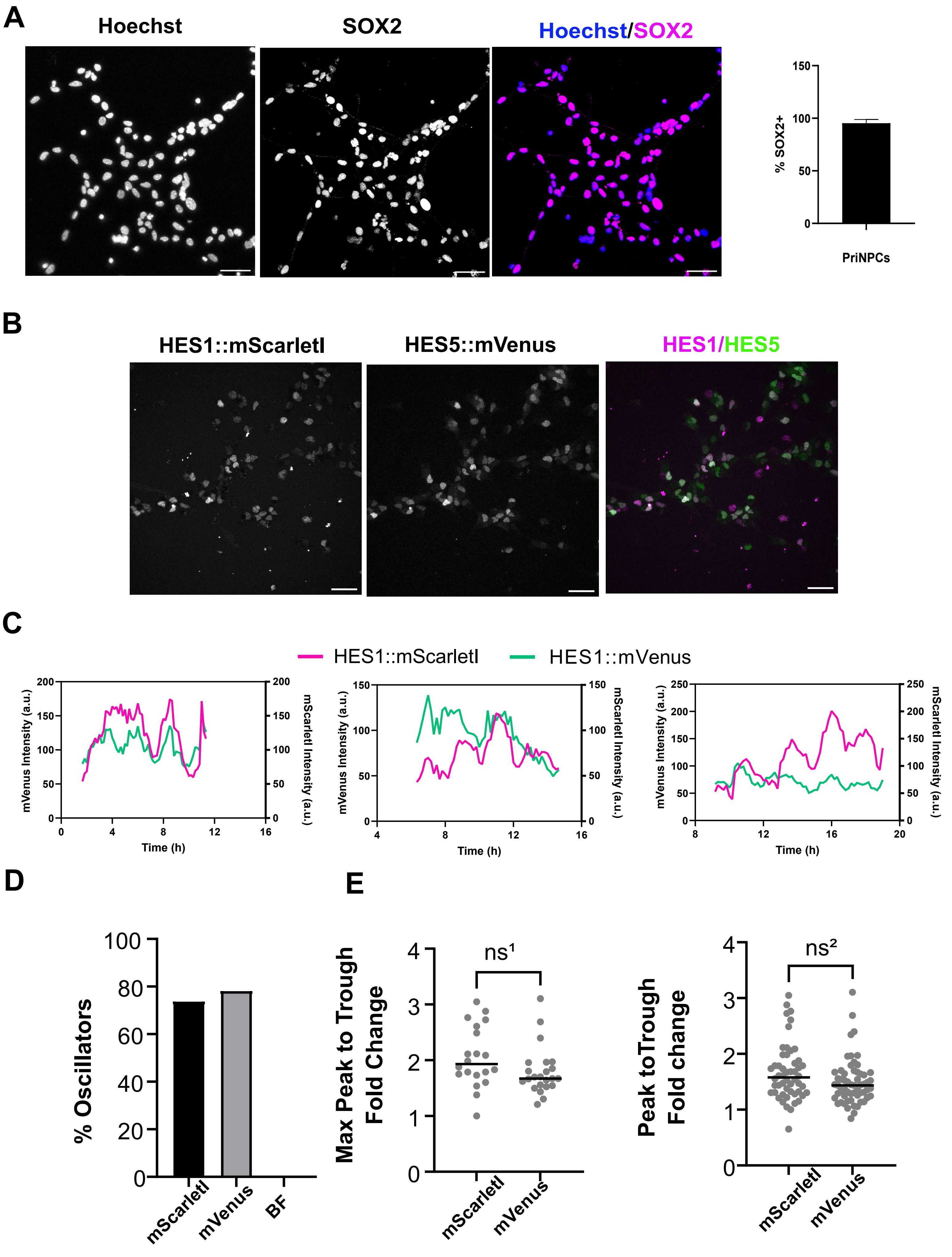

### Supplementary Figure 2

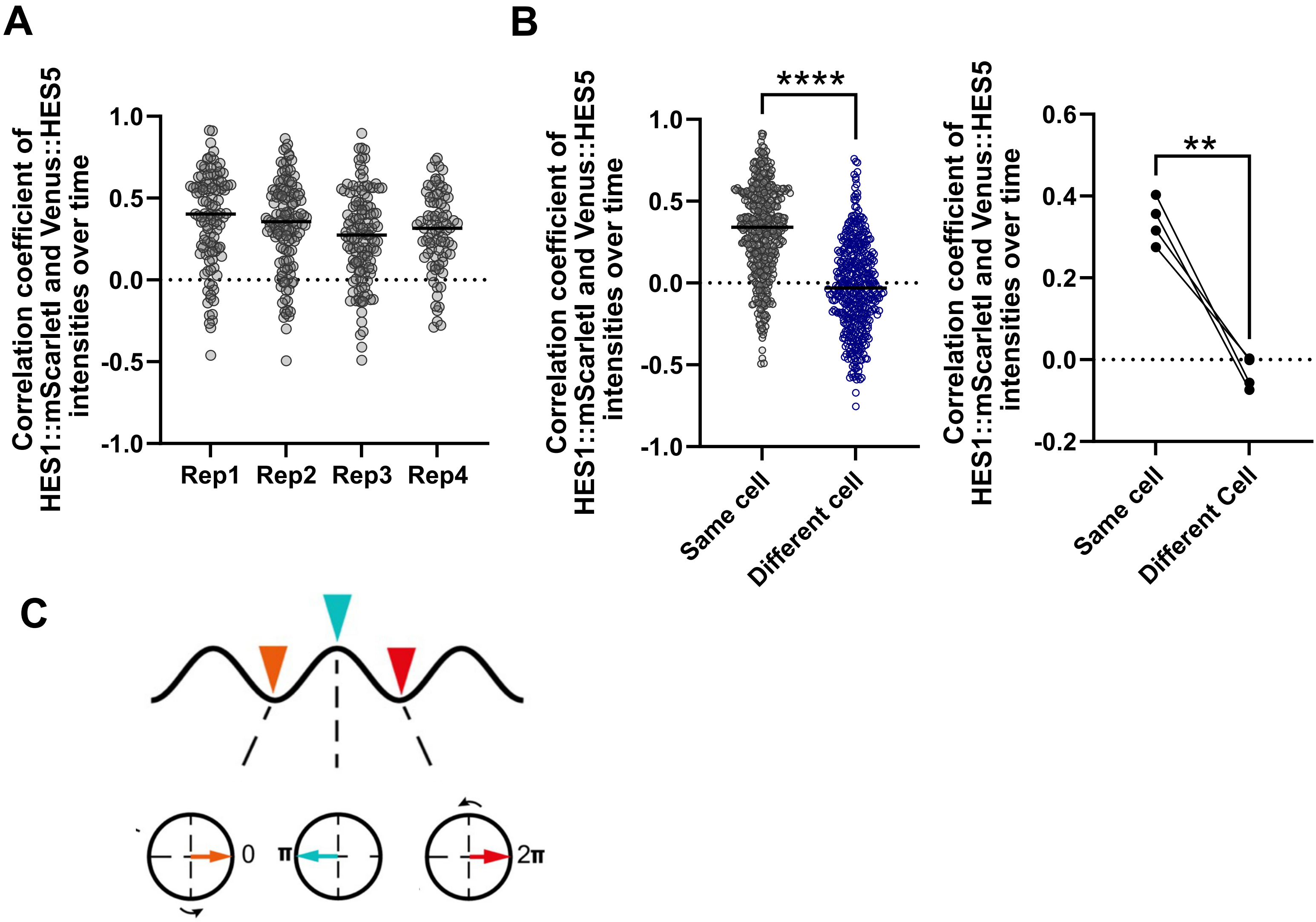

### Supplementary Figure 3

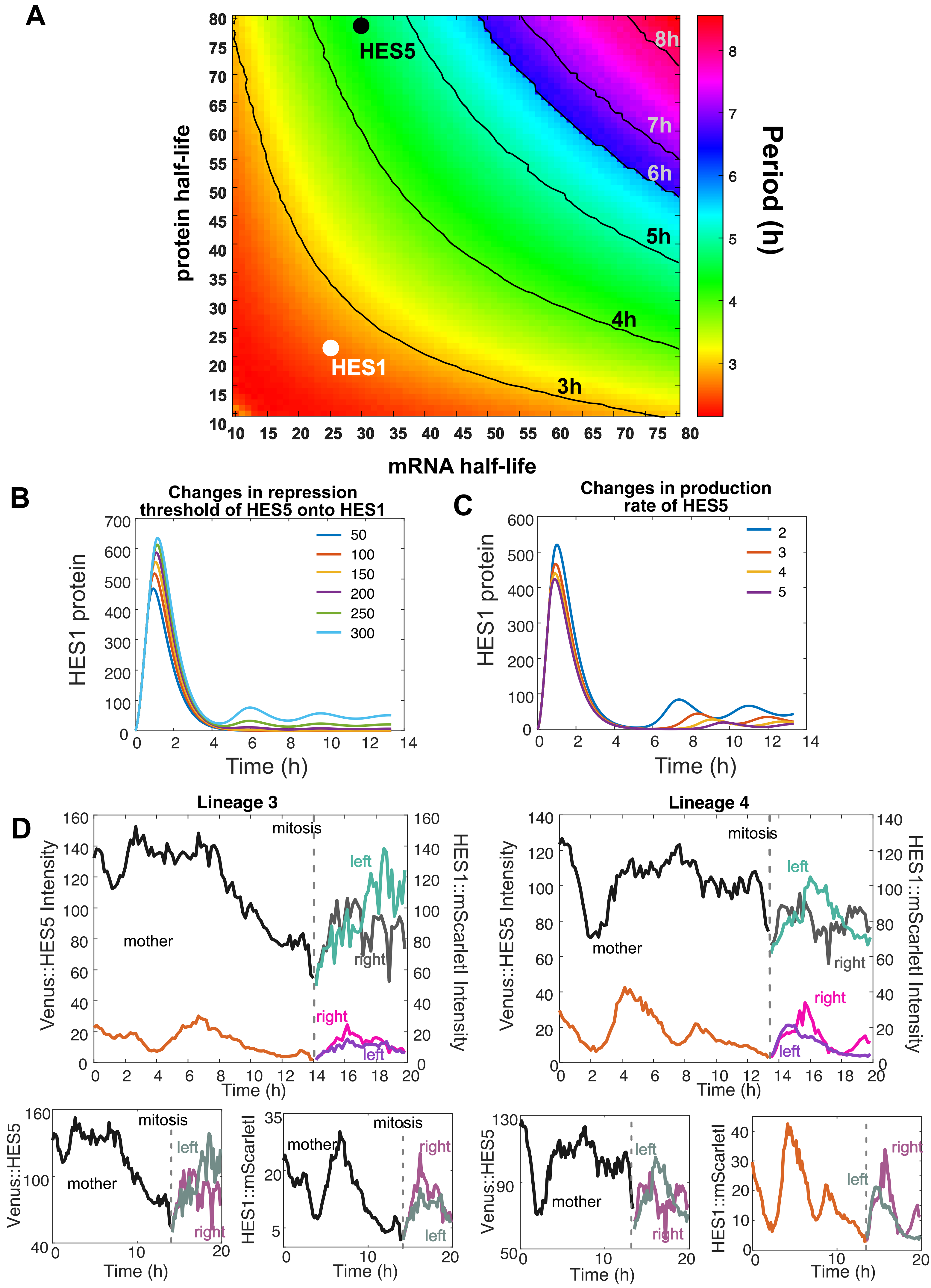

### Supplementary Figure 4

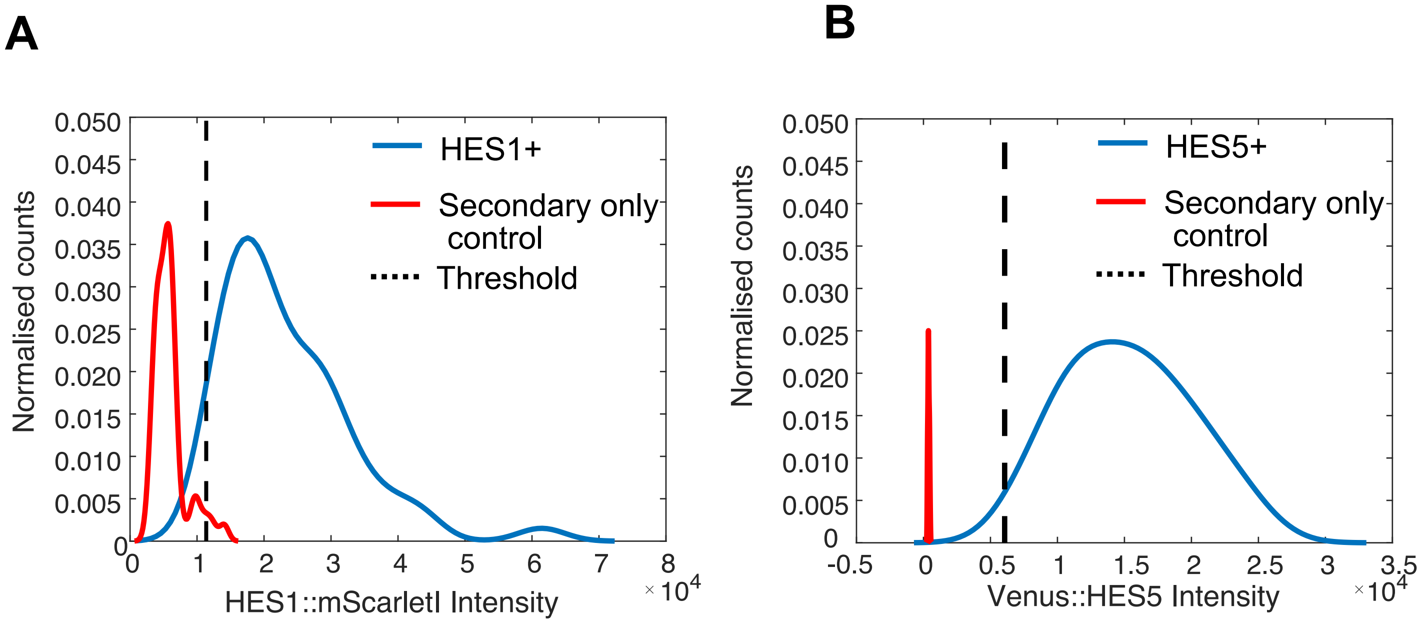

**Supplementary Figure 1. HES1 tagged with both mVenus and mScarlet-I shows similar fold-changes in oscillatory expression dynamics. Related to Figure 1 and 2.**

**(A)** SOX2 expression in primary neural progenitor cells detected by immunofluorescence; 40 x objective, scale bar is 40um; (right-most panel) percentage of SOX2+ nuclei as mean and SD from 3 independent experiments with a total of 1,075 nuclei analysed.

**(B)** Primary neural progenitor cell cultures containing a dual knock-in for HES1::mScarlet-I and HES1::mVenus; 40x objective, scale bar 40um.

**(C)** Representative examples of HES1::mScarletI and HES1::mVenus oscillatory intensity variations observed in the same nucleus over time.

**(D)** Percentage of oscillatory cells as detected from mScarlet-I or mVenus versus aperiodic variations in the bright field (BF) signal in the same nucleus; 1 independent experiment with 23 tracks analysed.

**(E)** Comparison of maximum peak to trough (left) and overall peak to trough values observed with mScarlet-I versus mVenus in the same cells; markers indicate peaks, line indicates median of 1 experiment; unpaired t-test, 2 tailed non-significant, $ns^1$=0.0873, $ns^2$=0.0536.

**Supplementary Figure 2. Correlation analysis of HES1 and HES5 expression in the same primary spinal cord neural progenitor cells. Related to Figure 3.**

**(A)** Pearsons correlation coefficient computed from HES1::mScarlet-I and Venus::HES5 timeseries observed in the same nucleus over time in 4 independent experiments with a total of 446 tracks; markers indicate single nuclei, bars indicate median per experiment.

**(B)** Pearsons correlation coefficient from data in (A) compared against correlation coefficient values obtained when cross-pairing HES1::mScarletI in one nucleus with Venus::HES5 in another nucleus selected at random; (left panel) markers indicate individual nuclei, lines indicate median of 4 pooled independent experiments, Mann-Whitney 2 -tailed test with $p<0.0001$; (right panel) markers indicate paired medians per experiment, paired t-test, 2-tailed with $p=0.0037$.

**(C)** Diagram depicting phase angle reconstruction from a wave resulting in values ranging from 0 to 2pi over the course of a complete oscillation cycle.

**Supplementary Figure 3. Mathematical model predicts HES1 and HES5 period based on experimental degradation rates. Related to Figures 4&5.**

**(A)** Period estimates (measured as average peak to peak intervals) from a free running model of HES at different values of mRNA and protein half-life; markers indicate known values reported for HES1 (white marker, mRNA half-life=25min (Bonev et al., 2012); protein half-life =22min (Hirata et al., 2002)) and HES5 (black marker, mRNA half-life=30min; protein half-life =80min (Manning et al., 2019)). Model parameters: $\mu_m = \mu_m = 1min^{-1}$; $P_0 = 390; n = 5;\ \tau = 29min.$

**(B)** Simulations of progressive HES5 repression onto HES1 through changes in cross-repression threshold values ($P_{051} \in [50,300]$). Model parameters: $P_{015} = P_{01} = P_{05} = 390$ , $\mu_{p1} = \frac{\ln(2)}{22}$ min$^{-1}$, $\mu_{m1} = \frac{\ln(2)}{25}$ min$^{-1}$; $\mu_{p5} = \frac{\ln(2)}{80}$ min$^{-1}$, $\mu_{m5} = \frac{\ln(2)}{25}$ min $^{-1}$, $\alpha_{m1} = \alpha_{p1} = \alpha_{m5} = \alpha_{p5} = 1$ min$^{-1}$; $\tau = 29$ min; $n = 5$.

**(C)** Simulations of progressive HES5 repression onto HES1 through changes production rates ($\alpha_{m5} = \alpha_{p5} \in [2,5]$). Model parameters: $P_{015} = P_{01} = P_{05} = 390$, $\mu_{p1} = \frac{\ln(2)}{22}$ min$^{-1}$, $\mu_{m1} = \frac{\ln(2)}{25}$ min$^{-1}$; $\mu_{p5} = \frac{\ln(2)}{80}$ min$^{-1}$, $\mu_{m5} = \frac{\ln(2)}{25}$ min $^{-1}$, $\alpha_{m1} = \alpha_{p1} = 1$ min$^{-1}$; $\tau = 29$ min; $n = 5$.

**(D)** Representative examples of HES1::mScarletI and Venus::HES5 fluorescent intensities observed in dividing mES-derived neural progenitor lineages comprising mother cell (division time unknown) and 2 daughter cells (left and right); intensities of mScarletI and Venus are not directly comparable however the same fluorophore can be compared between mother, left and right; lineages3&4 indicate that both left and right retain levels of HES1 and HES5 comparable to the mother cell following division; lineages 3&4 were collected in the same experiment as lineages 1&2 included in **Figure 5D.**

**Supplementary Figure 4. Quantification of HES1+ and HES5+ cells from tissue data. Related to Figure 6.**

**(A-B)** Intensity threshold for selecting HES1+ and HES5+ from data in **Figure 6B-F**; the distribution of nuclear intensity values for HES1:mScarletI detected by anti-RFP and Venus::HES5 detected by anti-GFP with intensity above threshold (dashed line) is compared against the distribution of nuclear intensities observed in slices with secondary only control (no anti-RFP, no anti-GFP).
